# The Aquarius/EMB-4 helicase licenses co-transcriptional gene silencing

**DOI:** 10.1101/089763

**Authors:** Alper Akay, Tomas Di Domenico, Kin M. Suen, Amena Nabih, Guillermo E. Parada, Mark Larance, Ragini Medhi, Ahmet C. Berkyurek, Xinlian Zhang, Christopher J. Wedeles, Ping Ma, Angus I. Lamond, Martin Hemberg, Julie M. Claycomb, Eric A. Miska

**Affiliations:** Gurdon Institute, University of Cambridge, Tennis Court Road, Cambridge, CB2 1QN, United Kingdom; Department of Genetics, University of Cambridge, Downing Street, Cambridge, CB2 3EH, United Kingdom; Wellcome Trust Sanger Institute, Wellcome Trust Genome Campus, Cambridge, CB10 1SA, United Kingdom; Department of Molecular Genetics, University of Toronto, Toronto, ON, M5S 1A8, Canada; Centre for Gene Regulation and Expression, School of Life Sciences, University of Dundee, Dundee, DD1 5EH, United Kingdom; Department of Statistics, University of Georgia, Athens, GA 30602, USA

## Abstract

Small RNAs (sRNAs) play an ancient role in genome defence against transposable elements. In animals, plants and fungi small RNAs guide Argonaute proteins to nascent RNA transcripts to induce co-transcriptional gene silencing. In animals the link between small RNA pathways and the transcriptional machinery remains unclear. Here we show that the *Caenorhabditis elegans* germline Argonaute HRDE-1 physically interacts with the conserved RNA helicase Aquarius/EMB-4. We demonstrate that the Aquarius/EMB-4 helicase activity is required to initiate small RNA-induced co-transcriptional gene silencing. HRDE-1 and Aquarius/EMB-4 are required to silence the transcription of overlapping sets of transposable elements. Surprisingly, removal of introns from a small RNA pathway target abolishes the requirement for Aquarius/EMB-4, but not HRDE-1, for gene silencing. We conclude that the Aquarius/EMB-4 helicase activity allows HRDE-1/sRNA complexes to efficiently engage nascent RNA transcripts - in competition with the general RNA processing machinery. We postulate that Aquarius/EMB-4 facilitates the surveillance of the nascent transcriptome to detect and silence transposable elements through small RNA pathways.

## INTRODUCTION

Transposable elements (TEs) are a feature of almost all eukaryotic genomes, which if left unchecked present a considerable danger to genome integrity and species survival. Therefore, organisms have evolved robust pathways to silence the expression of transposable elements and to restrict their mobility (Slotkin and Martienssen, 2007). Among these, DNA and chromatin modification and small RNA-mediated silencing are ancient and conserved mechanisms. Small RNA pathways in eukaryotes are all related to RNA interference mechanisms (RNAi) (Fire et al., 1998):21-32 nucleotide (nt) small RNAs (sRNAs) are bound by Argonaute superfamily proteins, interact with target RNAs through Watson-Crick base-pairing and initiate silencing of these targets. Such sRNA-mediated silencing can be post-transcriptional on mRNAs in the cytoplasm (PTGS) or co-transcriptional on nascent transcripts in the nucleus (coTGS). The latter provides the potential to couple sRNA-mediated silencing to DNA and chromatin-based gene regulatory pathways. The mechanisms by which such sRNA-mediated silencing affects DNA methylation and/or chromatin modification remain largely unknown, particularly in animals (Holoch and Moazed, 2015; Weick and Miska, 2014).

Most organisms have also evolved sRNA amplification systems to enhance sRNA-mediated silencing. The ancestral amplification mechanism is based on RNA-dependent RNA polymerases (RdRPs), which use target RNAs as templates to generate secondary sRNAs. Some animals, including mammals, have lost RdRPs but have instead evolved the Ping-pong amplification system (Aravin et al., 2008; Brennecke et al., 2007). The Ping-pong pathway amplifies Piwi-interacting RNAs (piRNAs), an animal-specific class of sRNAs, through double-stranded target RNA intermediates and Argonaute RNase activity (RNA slicing). piRNAs are named after a subfamily of Argonautes, the Piwi proteins. (Cox et al., 1998). Piwi proteins and piRNAs act in TE silencing in the germline of many animals. Another distinction of piRNAs is that they act *in trans*, where piRNAs generated from genomic clusters silence TEs throughout the genome. piRNA clusters can be thought of as a memory of TEs within the genome of an organism analogous to the guide RNA cluster/CRISPR systems of prokaryotes. Like other sRNAs, piRNAs have been shown to act through PTGS and coTGS mechanisms in nematodes, insects, fish and mice (Bagijn et al., 2012; Carmell et al., 2007; Kuramochi-Miyagawa et al., 2008; Le Thomas et al., 2013). *C. elegans* has both PTGS and coTGS mechanisms in the soma and the germline (Weick and Miska, 2014). In the germline sRNAs and piRNAs can initiate coTGS through a two-step mechanism. *C. elegans* piRNAs are 21 nt RNAs with a 5′ uracil (21U-RNAs) that are bound by the PRG-1 Piwi protein in the germline cytoplasm (Batista et al., 2008; Das et al., 2008; Wang and Reinke, 2008). Once a PRG-1/piRNA complex has recognized a target RNA it recruits an RdRP-containing complex to generate 22 nt antisense sRNAs with a 5′ guanine (22G-RNAs) (Bagijn et al., 2012; Pak and Fire, 2007).22G-RNAs are then bound by the Argonaute HRDE-1 and imported into the nucleus (Ashe et al., 2012; Buckley et al., 2012; Shirayama et al., 2012). A HRDE-1/22G-RNA complex is thought to directly interact with nascent transcripts. Genetic screens have identified several additional factors that are required for HRDE-1 mediated coTGS including NRDE-1, NRDE-2 and NRDE-4, the H3K9 histone methyltransferases SET-25, SET-32 and the heterochromatin protein 1 (HP1) homolog HPL-2 (Ashe et al., 2012; Buckley et al., 2012; Burkhart et al., 2011; Guang et al., 2008; Sapetschnig et al., 2015; Shirayama et al., 2012). However, how HRDE-1 links sRNA-mediated silencing to coTGS and chromatin modifications remains unknown.

Transcription and mobility of TEs is actively suppressed by this two-step piRNA/22G-RNA coTGS system in *C. elegans*. Interestingly, piRNA- and sRNA-mediated coTGS can last for multiple generations (Ashe et al., 2012; Luteijn et al., 2012; Shirayama et al., 2012),can be the source of non-genetic transgenerational effects (Ashe et al., 2012; Jobson et al., 2015; Ni et al., 2016; Rechavi et al., 2014) and loss of genes in the piRNA and the 22G-RNA pathway result in a multi-generational loss of fertility (de Albuquerque et al., 2015; Buckley et al., 2012; Phillips et al., 2015; Simon et al., 2014).

Eukaryotic mRNA transcription is a complex, multi-step process. First, the RNA polymerase II holoenzyme assembles on genomic DNA at a transcription start site. Second, nascent transcripts are produced by the elongating RNA polymerase II. Third, nascent RNA transcripts are processed to mature mRNAs through the assembly of multiple large ribonucleic acid protein (RNP) complexes to carry out 5′ capping (Gonatopoulos-Pournatzis and Cowling, 2014),splicing (Carrillo Oesterreich et al., 2016) and 3′ poly(A) tailing (Shatkin and Manley, 2000). All of these steps are required to protect transcripts from degradation and to ensure that mRNA is successfully exported into cytoplasm for protein translation. Of these co-transcriptional events, splicing is probably the most complex of all requiring more than 200 proteins and many non-coding RNAs (Nilsen, 2003; Wahl and Lührmann, 2015). Pre-mRNA splicing by the spliceosome is immediately followed by the assembly of a set of proteins known as the exon-junction complex (EJC). EJC functions in export, localisation and translation of mRNAs. Assembly of different EJC components can determine the fate of mRNAs and EJC can be considered a regulatory hub between transcription and translation (Le Hir et al., 2016).

Interestingly, most TEs are adapted to the host transcription and RNA processing machineries and exploit them for their own expression and mobility (Cowley and Oakey, 2013; Rebollo et al., 2012). Furthermore, TEs have been domesticated in such a way that their regulation can contribute to gene regulation (Cowley and Oakey, 2013; Rebollo et al., 2012).

Here we sought to gain further insight into coTGS mechanisms in animals. Specifically, we used a proteomic approach to identify protein interactors of the germline Argonaute HRDE-1 in *C. elegans*. We found that HRDE-1 interacts with components of the elongating RNA polymerase machinery including the spliceosome and the EJC. One HRDE-1 interacting factor is the conserved RNA helicase Aquarius/EMB-4, which binds introns and recruits the EJC to newly spliced transcripts. HRDE-1 and Aquarius/EMB-4 act to silence an overlapping set of TEs and TE-containing genes. The helicase activity of Aquarius/EMB-4 is required for HRDE-1/sRNA-mediated coTGS. Surprisingly, removal of introns of a coTGS target removes the requirement for Aquarius/EMB-4 in sRNA-mediated silencing. Thus Aquarius/EMB-4 activity allows HRDE-1 access to the nascent RNA transcript. In summary, Aquarius/EMB-4 enables efficient sRNA-mediated genome surveillance by licensing coTGS during pre-mRNA processing.

## RESULTS

### SILAC proteomics identifies protein interactors of the germline nuclear Argonaute HRDE-1

To further our understanding of coTGS in animals and as a complement to previous genetic approaches we sought to identify proteins interacting with the nuclear Argonaute HRDE-1 by protein immunoprecipitation (IP) coupled to SILAC (Stable isotope labelling of amino acids in cell culture) proteomics (Ong et al., 2002),using the SILAC labeling for nematodes approach (Akay et al., 2013; Fredens et al., 2011; Larance et al., 2011) (Figure 1A). We produced whole-cell protein extracts from wild-type young adult *C. elegans* grown on “heavy” isotopes and *hrde-1(tm1200)* mutant animals grown on “light” isotopes. We chose the young adult stage as the germline is then fully developed and endogenous HRDE-1 expression is at its peak (Ashe et al., 2012). We then immunoprecipitated a 1:1 mix of protein extracts of both genotypes using an anti-HRDE-1 antibody followed by liquid chromatography tandem-mass spectrometry (LC/MS-MS). Based on three biological replicates LC/MS-MS identified 130 candidate HRDE-1 interacting proteins. For further analysis and to take advantage of prior proteomics data from mammals, we focused on 53 proteins with human orthologs that are also classified as being nuclear (Uhlén et al., 2015; UniProt Consortium, 2015). We used the STRING database (Szklarczyk et al., 2015) to identify known protein complexes between these factors. 28 of these 53 proteins were connected with each other based on known interactions and the majority of the proteins represent RNA processing factors, RNA polymerase II subunits and nuclear pore complex components (Figure 1B, Figure S1A). In the fruitfly *Drosophila melanogaster* the Piwi protein has an analogous function to HRDE-1 in coTGS in the germline (Weick and Miska, 2014). Interestingly, eight proteins identified in our HRDE-1 IPs were also found in *D. melanogaster* PIWI IPs. In addition, *D. melanogaster* orthologs of 13 of our HRDE-1 interactors are required for the regulation of transposable elements when assayed by *in vivo* RNAi (Figure 1C, Table S1) (Czech et al., 2013; Le Thomas et al., 2013). Overall, our SILAC based HRDE-1 proteomics identified a set of conserved proteins that might bridge sRNA pathways, RNA processing and TE silencing.

**Figure 1.**
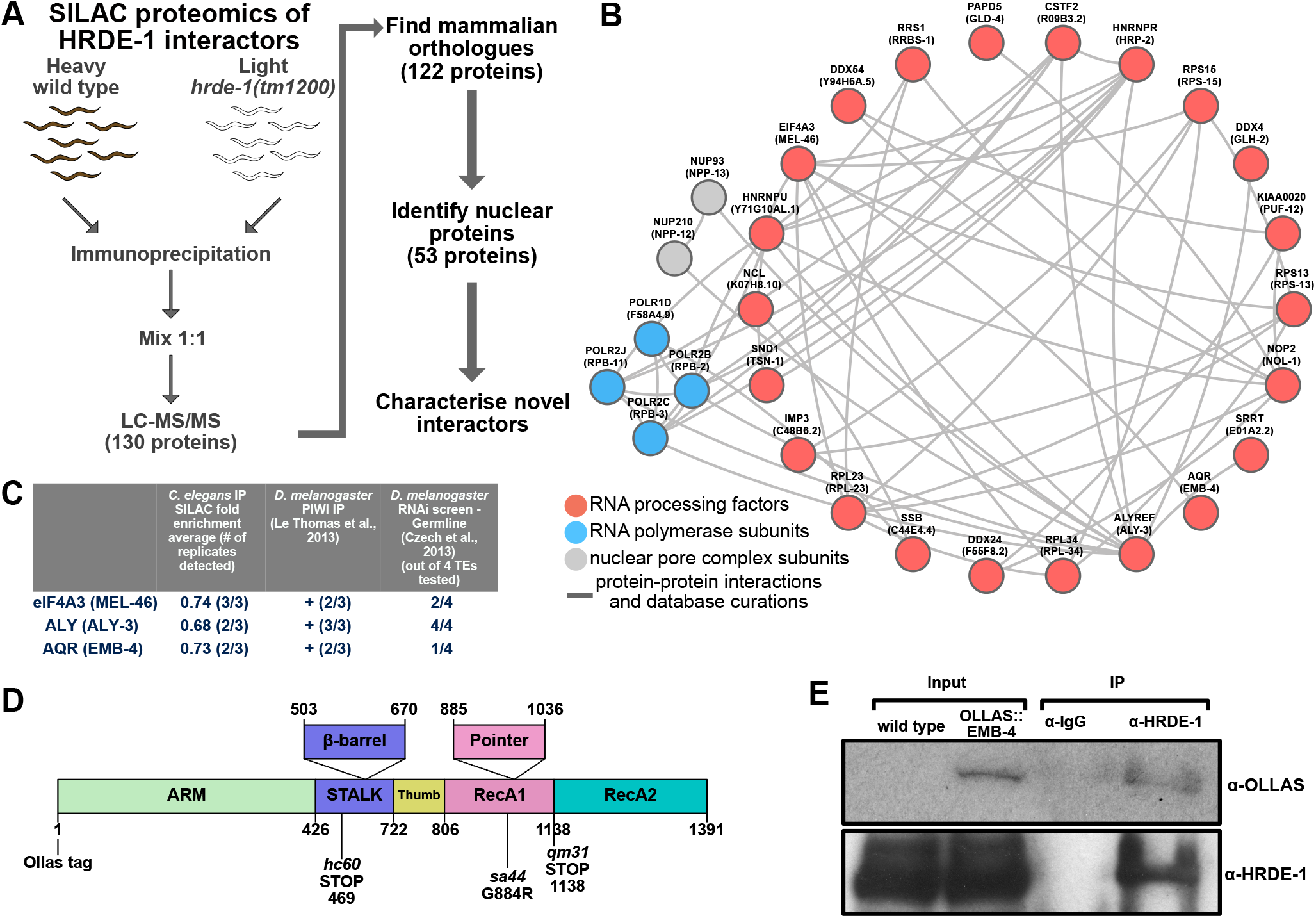
SILAC proteomics identifies Aquarius/EMB-4 as an interactor of the nuclear Argonaute HRDE-1 (A) SILAC labelling and immunoprecipitation scheme for wild type (heavy labelled) and *hrde-1(tm1200)* mutants (light labelled) using anti-HRDE-1 antibodies. (B) Known protein interactions identified using the STRING database for mammalian orthologues of nuclear HRDE-1 interactors (Figure 1A). Grey lines indicate known protein-protein interactions. Red circles highlight known RNA processing factors, blue circles highlight RNA pol II subunits and grey circles highlight nuclear pore complex subunits. (C) Aquarius and two exon-junction complex proteins eIF4A3 and ALY are detected in HRDE-1 IPs (second column, mean log2 fold enrichment heavy / light), detected in *D. melanogaster* PIWI IPs (third column, number of IPs detected/total number of IPs) and effect on TE de-silencing in *D. melanogaster* upon RNAi knock-down (fourth column, number of TEs de-silenced out of four tested). (D) EMB-4 domain structure based on the mammalian homolog Aquarius and the position of *emb-4* mutations used in this study. (E) Validation of protein-protein interaction between HRDE-1 and EMB-4 using anti-HRDE-1 antibodies to immunoprecipitate HRDE-1 complexes in CRISPR tagged Ollas-EMB-4 strain.

### The Aquarius RNA helicase ortholog EMB-4 is nuclear, germline-enriched and interacts with HRDE-1

Of the candidate HRDE-1 protein interactors we were particularly curious about EMB-4 as its interaction with nuclear Argonaute proteins appeared to be conserved in *D. melanogaster* (Figure 1C). EMB-4 is the *C. elegans* ortholog of Aquarius (AQR), a large scaffolding protein that includes two RecA helicase domains (Figure 1D). Aquarius/EMB-4 is conserved in all eukaryotes with an intact RNAi pathway, including the fission yeast *Schizosaccharomyces pombe*, but not in *Saccharomyces cerevisiae*, which lacks RNAi (Drinnenberg et al., 2011) (Figure S1B,C). As Aquarius has a conserved role in nascent RNA processing in eukaryotes (De et al., 2015; Wahl and Lührmann, 2015) we wondered if Aquarius/EMB-4 might provide new insights into the interface between the nuclear RNAi pathway and the general RNA processing machinery. Therefore we validated the interaction between HRDE-1 and EMB-4 independently. First we generated an N-terminal epitope-tagged version of EMB-4 *in vivo* using CRISPR/Cas9 genome engineering (Paix et al., 2015) (Ollas tag, Figure 1D). We then tested the interaction between HRDE-1 and EMB-4-Ollas using an anti-Ollas (Park et al., 2008) and an anti-HRDE-1 (Ashe et al., 2012) antibody through IP followed by western blotting. As shown in Figure 1E we found that EMB-4-Ollas interacts specifically with HRDE-1 in whole-cell extracts. In addition, we validated this interaction using an anti-EMB-4 antibody and a Flag-tagged version of HRDE-1 generated by MosSCI transgenesis (Frøkjær-Jensen et al., 2008; Shirayama et al., 2012) (Figure 1D, Figure S1E,F). Next we asked whether EMB-4 is co-expressed with HRDE-1 *in vivo*. We found that *emb-4* mRNA was highly enriched in the germline, the embryo and early larval stages of *C. elegans* (Figure 2A). Using an anti-EMB-4 antibody we observed the same for the endogenous EMB-4 protein (Figure 2B). Taking advantage of temperature-sensitive germline mutants of *C. elegans* we found that in gravid adult animals the majority of EMB-4 mRNA and protein is germline restricted (Figure 2A,B). This is of particular interest as HRDE-1 expression is germline specific (Ashe et al., 2012). Within the germline we find that EMB-4 is highly enriched in germline nuclei and closely associated with chromatin in the mitotic, transition and pachytene region of the germline as well as in oocytes (Figure 2C, Figure S2). We conclude that EMB-4 is expressed in germline nuclei and is a *bona fide* HRDE-1 interacting protein.

**Figure 2.**
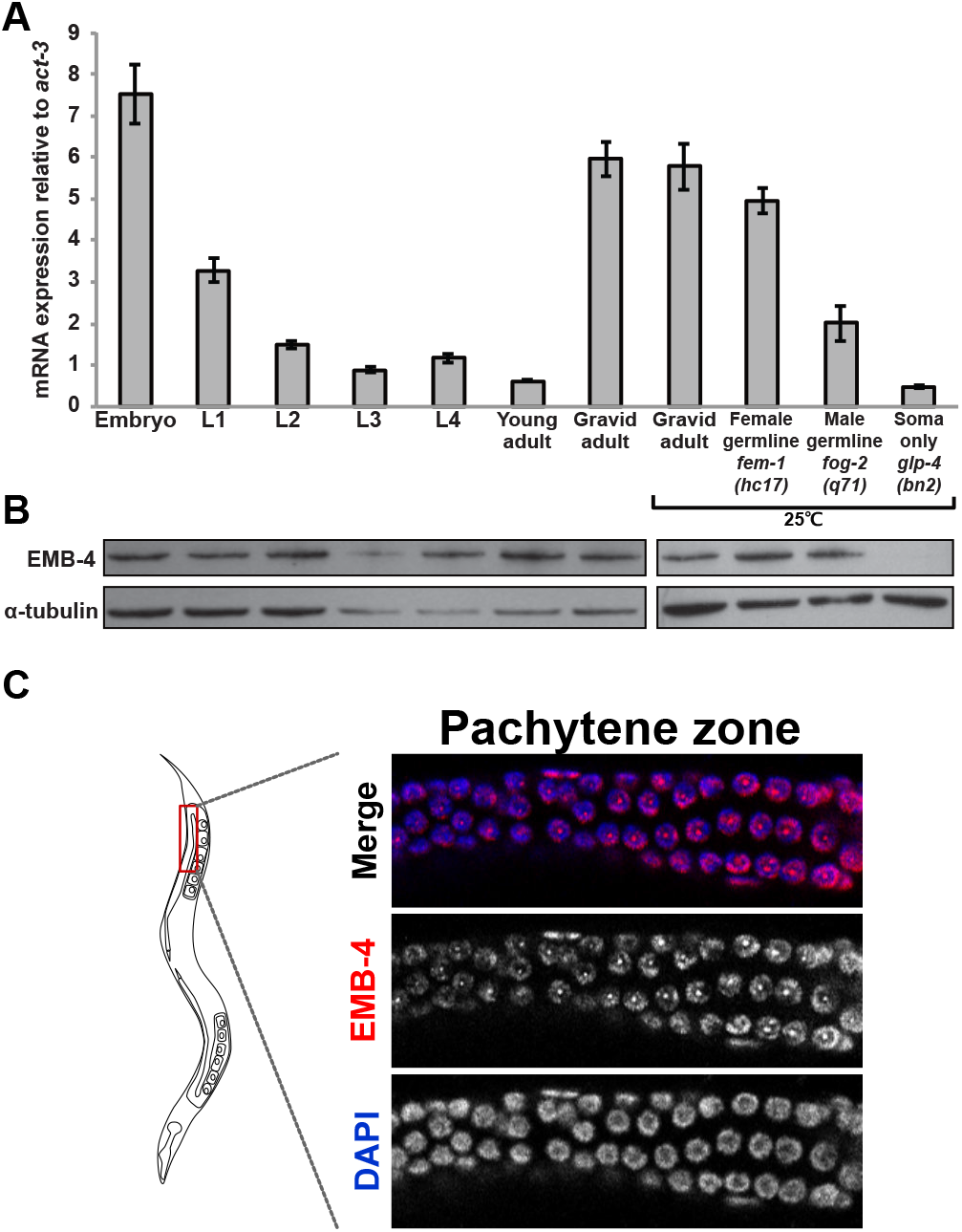
Aquarius/EMB-4 is highly enriched in germ cell nuclei. (A) qRT-PCR analysis of *emb-4* mRNA expression at different developmental stages (embryo - gravid adult), in animals lacking sperm (female germline *fem-1(hc17*)), in animals lacking oocytes (male germline *fog-2(q71)*) and in animals lacking a germline (soma only *glp-4(bn2)*).*fem-1(hc17), fog-2(q71)* and *glp-4(bn2)* are temperature sensitive mutants and were grown at 25℃ alongside the wild type gravid adult control). (B) Western blot analysis of EMB-4 protein levels using anti-EMB-4 antibodies for the same conditions as in Figure 2A. (C) Localisation of EMB-4 protein in the germline of adult animals.

### Aquarius/EMB-4 is required for piRNA-mediated co-transcriptional gene silencing

Next, we examined if, like the Argonaute HRDE-1, Aquarius/EMB-4 is required for coTGS in the germline. For this purpose, we used a piRNA sensor strain we generated previously (Bagijn et al., 2012). The piRNA sensor (*mjIs144)* is a transgene that drives the expression of a GFP histone H2B fusion protein through the germline-specific *mex-5* gene promoter and is inserted as a single-copy on chromosome II. In addition, the piRNA sensor contains a 21 nt sequence perfectly complementary to the endogenous piRNA 21UR-1 (Figure 3A). In otherwise wild-type animals, the piRNA sensor is silenced by piRNA-mediated coTGS. However, mutations in the piRNA or germline nuclear RNAi pathway *e.g. prg-1* (Bagijn et al., 2012)*,prde-1* (Weick et al., 2014) or *hrde-1* (Ashe et al., 2012) lead to re-activation of the piRNA sensor. Taking advantage of two previously isolated alleles of *emb-4* (Checchi and Kelly, 2006; Katic et al., 2006),we tested the requirement of EMB-4 in piRNA-mediated gene silencing. We observed that two null mutants of *emb-4* have de-silenced the piRNA sensor in the germline of *C. elegans* as indicated by GFP expression in germline nuclei, as did *hrde-1* as the positive control while the piRNA sensor is silenced in otherwise wild-type animals (Figure 3B-C, S3A). We also quantified mRNA levels of piRNA sensor expression in these mutant backgrounds, which confirmed the de-silencing in *emb-4* and *hrde-1* mutant backgrounds (Figure 3C). Finally, we carried out an analogous experiment using an independent transgene targeted by the piRNA pathway (*ccSi1504*, gift from C. Frøkjær-Jensen). *ccSi1504* is integrated on chromosome V, expressed under *smu-1* promoter and utilises *smu-1* introns and contains SV40 and EGL-13 nuclear localisation signals. Both *emb-4* and *hrde-1* mutants de-silenced the *ccSi1504* transgene similar to the *mjIs144* transgene (Figure S3B). Our results demonstrate that EMB-4 is an essential factor for piRNA-mediated coTGS in the germline.

**Figure 3.**
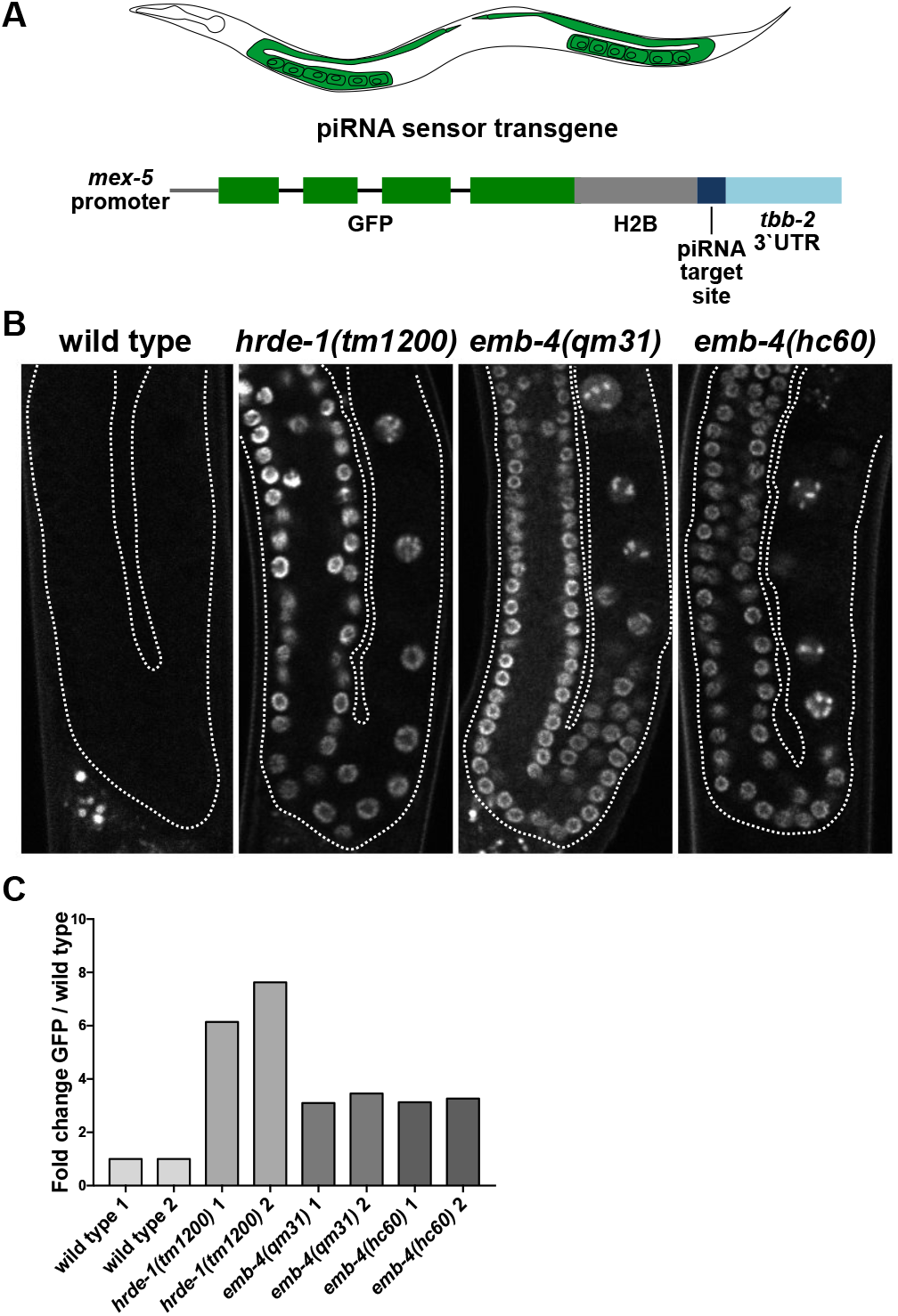
Aquarius/EMB-4 is required for the piRNA mediated silencing of a sensor transgene. (A) piRNA sensor transgene and its expression pattern in *C. elegans* germline (B) Fluorescent microscope images of wild type and mutant animal germlines with an integrated single copy piRNA sensor transgene (germline boundaries are marked up by white dotted lines). (C) qRT-PCR analysis of GFP expression of the adult animals as in Figure 3B (two biological replicates with at least two technical replicates).

### A functional RNA helicase domain of Aquarius/EMB-4 is required to establish co-transcriptional gene silencing

The molecular function of Aquarius/EMB-4 in vivo remains unknown in any organism. However, previous studies identified Aquarius as intron binding and associated with spliceosome and exon-junction complex recruitment. In vitro experiments suggested a role for the ATPase activity of the RecA1 helicase domain of human Aquarius/AQR in RNA unwinding and spliceosomal complex assembly (De et al., 2015). We noted that the previously isolated allele *sa44* of Aquarius/EMB-4 in *C. elegans* is a missense mutation that induces a G884R substitution in the RecA1 helicase domain (Katic et al., 2006). *emb-4(sa44)* was identified in a genetic suppressor screen and lacks the embryonic lethality associated with null mutants of *emb-4* (Checchi and Kelly, 2006; Katic et al., 2006). The G884R substitution is located within the loop region of the RecA1 helicase domain of EMB-4 (Figure 4A). Based on the structural alignment between human Aquarius/AQR, UPF1 and *C. elegans* EMB-4 (Figure S4A-D), G884R likely affects RNA-binding and thus helicase activity (Figure 4A) (Chakrabarti et al., 2011; Cheng et al., 2007; De et al., 2015). However, *emb-4(sa44)* mutants have wild-type Aquarius/EMB-4 protein levels (Figure S4E). Interestingly, *emb-4(sa44)* animals de-silence the piRNA sensor similar to *emb-4(null)* mutations (Figure 4B). We conclude that Aquarius/EMB-4 helicase activity is required for co-transcriptional gene silencing. We next asked whether Aquarius/EMB-4 is required for the establishment and/or the maintenance of co-transcriptional gene silencing. Taking advantage of the *emb-4(sa44)* mutants we addressed this using genetic crosses (Figure 4C,D). When *hrde-1(tm1200);mjIs144*(piRNA sensor) animals with a de-silenced piRNA sensor are crossed to *mjIs144* animals generating F1 animals heterozygous for *hrde-1*, piRNA sensor silencing is restored (Figure 4C). However, when carrying out an analogous cross of *hrde-1(tm1200);mjIs144;emb-4(sa44)* and *mjIs144;emb-4(sa44)* animals piRNA sensor silencing is not restored (Figure 4D). We conclude that Aquarius/EMB-4 is required for the establishment of co-transcriptional gene silencing.

**Figure 4.**
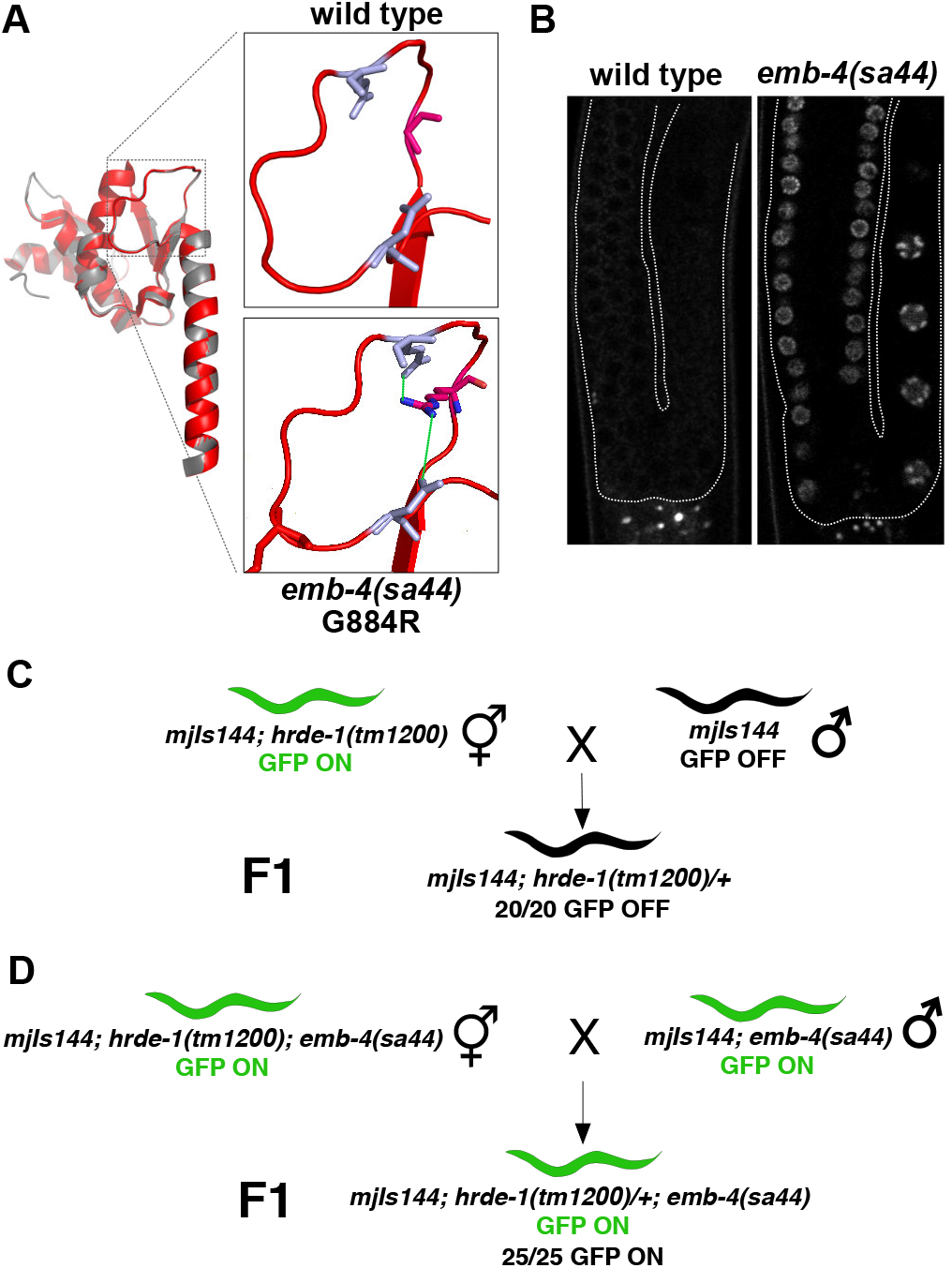
RNA helicase domain of Aquarius/EMB-4 is required for the establishment of transcriptional gene silencing. (A) Structural alignment of the RecA1 and pointer domains of human Aquarius (white) with *C. elegans* EMB-4 (red). Blown-up region showing the location of the G884R substitution found in *emb-4(sa44)* strain in the loop region of the helicase domain which forms the interface with the pointer domain that is embedded in the RecA1 helicase domain. Green lines show possible salt bridge interactions between amino acids. (B) Fluorescent images of wild type and *emb-4(sa44)* germlines with integrated piRNA sensor transgene (germline boundaries are marked up by white dotted lines). (C-D) Scheme of genetic crosses showing the effect of *emb-4(sa44)* mutation during the establishment of gene silencing. (C) In the control cross, a wild type copy of *hrde-1* (+) in the F1 heterozygous animals re-establishes complete piRNA sensor silencing. (D) Wild type copy of *hrde-1* fails to establish piRNA sensor silencing in the *emb-4(sa44)* homozygous background (number of F1 progeny is indicated below each cross).

### Aquarius/EMB-4 is required to silence transposable elements

Next we aimed to understand the role of Aquarius/EMB-4 on the endogenous transcriptome. We generated total RNA and small RNA expression profiles using high-throughput sequencing of wild-type animals and *hrde-1* and *emb-4* mutant animals (biological triplicates, deposited at ArrayExpress under E-MTAB-4877, see Experimental Procedures). First we took advantage of the fact that total RNA sequencing contains intronic sequence reads in addition to exonic sequence reads. Considering intronic reads as a measure of nascent RNA transcription, one can therefore infer changes of transcription rate in addition to steady-state mRNA levels. Such a method was recently reported as exon-intron split analysis (EISA) (Figure 5A) (Gaidatzis et al., 2015). Plotting the logarithm of the ratio of intronic reads from wild-type and mutant samples against the logarithm of the ratio of exonic reads of the same datasets one can infer the type of gene expression change between the samples; i.e. genes that vary in their transcription rate are aligned on the diagonal (yellow, Figure 5A-C), positive post-transcriptional regulation is above the diagonal (blue, Figure 5A-C) and negative post-transcriptional regulation below (grey, Figure 5A-C). Comparing three biological replicates each of wild-type animals, *hrde-1* and *emb-4* mutants (three replicates each of the *qm31* and *hc60* alleles, combined) we found that the majority of gene expression changes in *hrde-1* and *emb-4* mutants are due to changes in transcription rate (Figure 5B-C, % transcriptional changes: 83% in *hrde-1* and 73% in *emb-4*, transcriptionally up/down: 100/16 (*emb-4*), 34/5 (*hrde-1*)). Similarly, the piRNA sensor showed reduced transcription rates in both *hrde-1* and *emb-4* mutants (Figure 5B-C, green dot). These findings are consistent with the known role of HRDE-1 in co-transcriptional gene silencing (Ashe et al., 2012; Buckley et al., 2012). We conclude that EMB-4 similarly affects transcription rate, which is consistent with a model of EMB-4 and HRDE-1 acting together in co-transcriptional gene silencing. In addition, we conclude that EMB-4 has no major impact on pre-mRNA splicing as only a few genes showed a relative increase in intronic reads (Figure 5C, grey dots).

**Figure 5.**
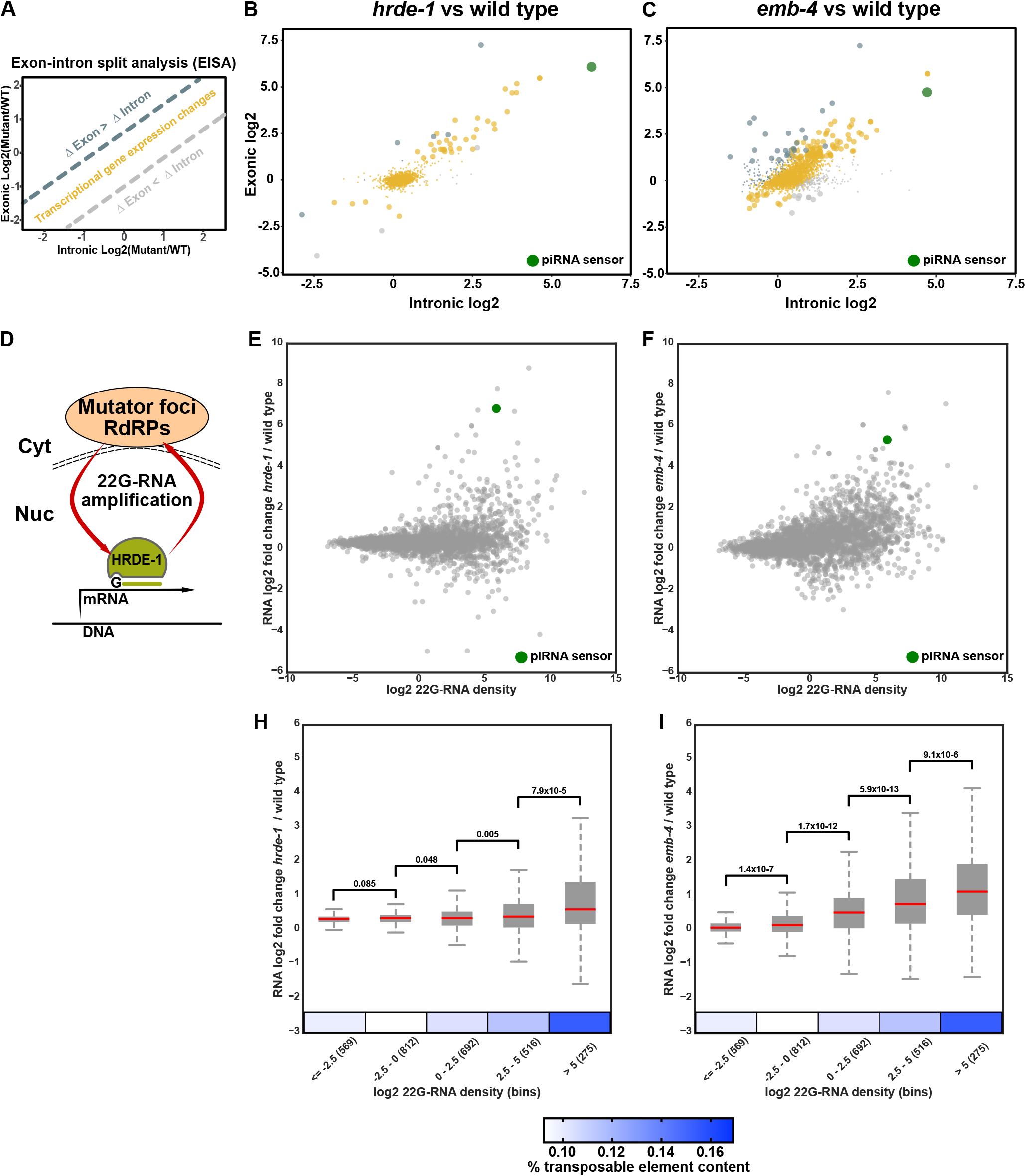
EMB-4 and HRDE-1 are required for transcriptional silencing of genes and transposable elements. (A) Exon-intron split analysis (EISA) for comparison of transcriptional and post-transcriptional gene expression changes. (B,C) Transcriptional and post-transcriptional gene expression changes are coloured as in (A). Significant gene expression changes are highlighted by larger dot size (large dots=mRNA log2FC ≥ 1, p-value ≤ 0.05). piRNA sensor transgene is highlighted by green. (D) Model showing 22G-RNA amplification in mutator foci which requires HRDE-1 for 22G-RNA transport and stability. (E, F) Comparison of RNA log2 fold change in *hrde-1* and *emb-4* mutants to log2 22G-RNA density in wild type animals (22G-RNA density=22G-RNAs in HRDE-1 IP in wild type / RNA RPKM in wild type). piRNA sensor transgene is highlighted in green. (H, I) Genes in (E, F) are grouped into bins of increasing 22G-RNA density as shown on X-axis (number of genes in each bin is shown in parentheses, boxes show the lower and upper quartiles, red line shows the median and *p-values* of two-sample *t-test* are shown above the box plots). Heat-map shows % abundance of transposable elements in each 22G-RNA bin.

Next we considered small RNA expression alongside total RNA expression to focus on the direct targets of co-transcriptional gene silencing. HRDE-1 bound sRNAs are 22G-RNAs, antisense to their target RNAs and generated by RdRPs at perinuclear foci called mutator bodies (Figure 5D). HRDE-1 is required for the stability and amplification of 22G-RNAs in the germline (Ashe et al., 2012; Buckley et al., 2012; Sapetschnig et al., 2015). We therefore calculated a 22G-RNA density for HRDE-1 targets as the ratio of HRDE-1 bound 22G-RNAs in the wild type and the expression levels of their target RNAs (Gerson-Gurwitz et al., 2016; Sapetschnig et al., 2015). We found that genes with high 22G-RNA density in wild-type animals, tended to be up-regulated in both *hrde-1* and *emb-4* mutants (Figure 5E-F).

For further analysis we grouped all genes into bins of increasing 22G-RNA density (bins 1-5, 275-569 genes per bin) and found a number of correlating features: HRDE-1 bound 22G-RNAs are enriched in germline 22G-RNA targets and depleted from somatic 22G-RNA targets as expected (Figure S5) (Gu et al., 2009a). In addition, 22G-RNA density correlates positively with piRNA targets and targets of the WAGO-1 pathway, which is known to overlap with HRDE-1 targets (Figure S5) (Lee et al., 2012). Finally, ERGO-1, ALG-3/4 and CSR-1 Argonautes are all required for additional endogenous RNAi pathways that also produce 22G-RNAs (Claycomb et al., 2009; Conine et al., 2010; Vasale et al., 2010). We found that HRDE-1 bound 22G-RNAs were overall depleted from ERGO-1 and ALG-3/4 targets and the density of HRDE-1 bound 22G-RNAs inversely correlated with CSR-1 targets (Figure S5). HRDE-1 bound 22G-RNAs are known to target transposable elements (Ni et al., 2014). Indeed, genes with the highest 22G-RNA density also had the highest TE content and showed the strongest mRNA up-regulation (Figure 5H-I) both in *hrde-1* and *emb-4* mutant animals. We therefore asked if EMB-4, like HRDE-1, regulates transposable element expression. Indeed, when grouping TEs into families, we found that several DNA transposons and retro-elements are over-expressed in *hrde-1* and/or *emb-4* mutants (Figure 6A). For instance, the CER-9 LTR is strongly induced in both *hrde-1* and *emb-4* mutants. Furthermore, CER-9 LTR induction is accompanied by complete loss of 22G-RNAs to the same locus in both mutant backgrounds (Figure 6B). While there is an overall good correlation in gene expression changes in *hrde-1* and *emb-4* mutants, some changes are unique to the individual mutants. For example, *bath-45* is a piRNA pathway target gene with multiple piRNA target sites on all of its exons. *bath-45* mRNA levels were upregulated in *hrde-1* but not in *emb-4* mutant animals (Figure S6A). On the contrary, two TEs, CER-15 LTR and Mirage1 showed strong upregulation only in *emb-4* mutants (Figure S6B-C). Altogether we conclude that HRDE-1 and Aquarius/EMB-4 act together to silence many endogenous 22G-RNA target loci, genes and transposable elements.

**Figure 6.**
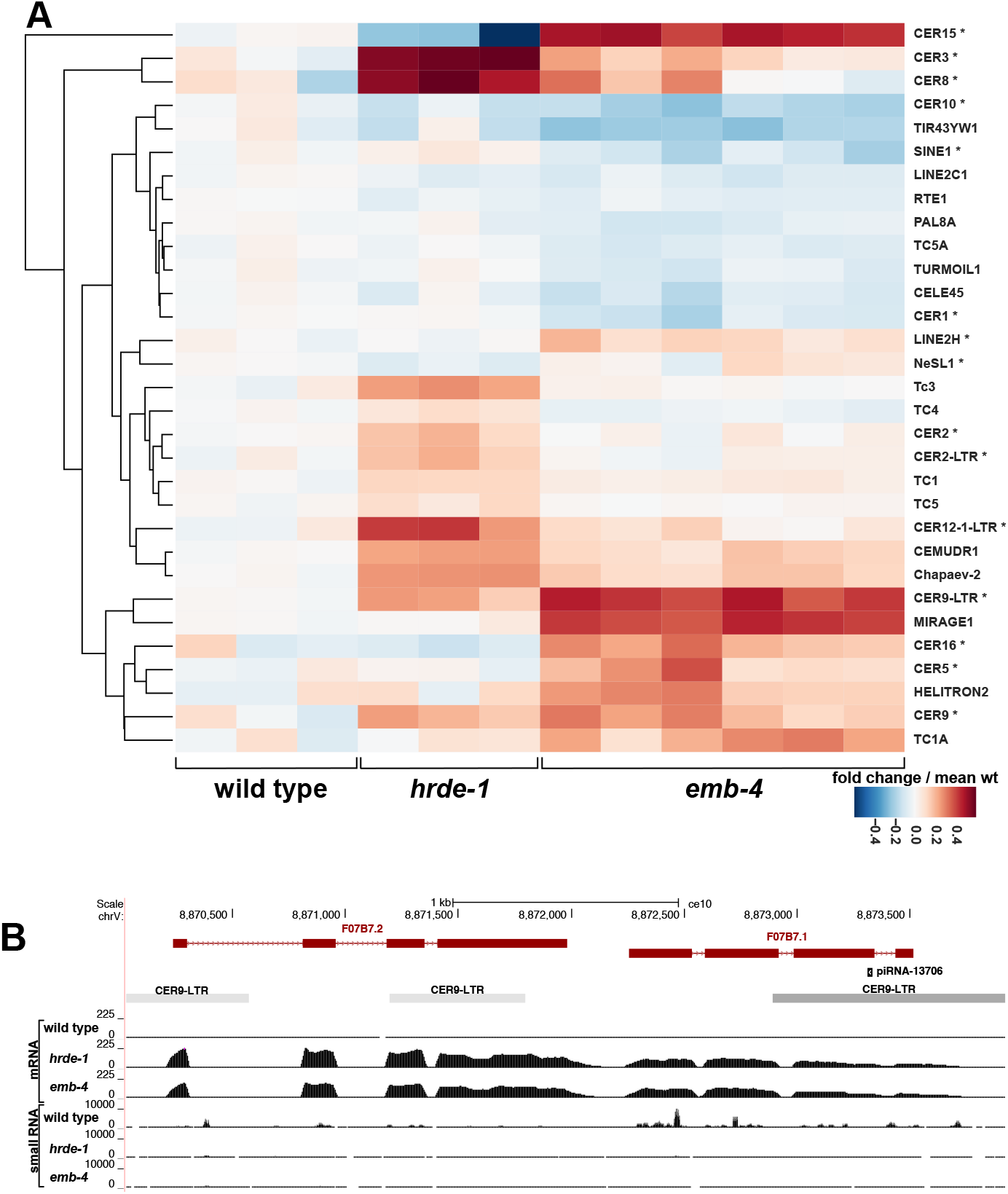
HRDE-1 and EMB-4 are required for the suppression of multiple transposable element families (A) Heat-map showing RNA fold change of transposable element families in wild type, *hrde-1* and *emb-4* mutants compared to mean wild type levels. (* indicates retro-element families) (B) CER-9 LTR region showing increased mRNA expression in *hrde-1* and *emb-4* mutants and reduced 22G-RNA reads.

### Aquarius/EMB-4 is specifically required to silence transcripts with introns

As Aquarius is known to bind introns and is required for RNP re-modelling during spliceosome and EJC assembly, we wondered if Aquarius/EMB-4 is specifically required to allow co-transcriptional gene silencing on nascent transcripts undergoing splicing. To test this we took advantage of the piRNA sensor, which requires HRDE-1 and EMB-4 for co-transcriptional gene silencing (Figures 3 and 5). We previously characterized 22G-RNA populations for the piRNA sensor in detail (Sapetschnig et al., 2015):upon initial piRNA targeting of the sensor transgene in the 3’UTR (Figure 3A), 22G-RNAs are generated proximal to the piRNA target site that are independent of HRDE-1 and the nuclear RNAi machinery and not sufficient for piRNA silencing. Subsequently, HRDE-1-dependent 22G-RNAs spread along the whole length of the transcript, including the *gfp* coding region, to induce co-transcriptional gene silencing (Figure S7A). We find that loss of HRDE-1 or Aquarius/EMB-4 resulted in loss of the majority of 22G-RNAs mapping to the coding region of the piRNA sensor, consistent with HRDE-1 and Aquarius/EMB-4 acting together in nuclear RNAi (Figure S7A). The piRNA sensor transgene (*mjIs144*) has three synthetic introns within the *gfp* gene sequence. These same introns are generally used by the community to promote efficient transgene expression in *C. elegans*. To test if Aquarius/EMB-4 is specifically required for co-transcriptional gene silencing of transcripts undergoing splicing we removed two of the three introns from the piRNA sensor transgene (*mjIs144*) to generate a new single intron piRNA sensor integrated into the same chromosomal location (*mjIs588*). We retained the single remaining intron, as an intronless transgene is unlikely to be expressed in *C. elegans* (Fire et al., 1990). As expected, the single intron piRNA sensor (*mjIs588*) was completely silenced in wild-type animals and fully de-silenced in *hrde-1* mutant animals (Figure 7A-B). In contrast, while the three-intron piRNA sensor required Aquarius/EMB-4 for silencing, the single intron piRNA sensor did not (Figure 7A-B). Importantly, GFP expression levels of the three intron piRNA sensor and the single intron piRNA sensor were similar in *hrde-1* mutants, confirming that the difference in Aquarius/EMB-4 dependence was not simply due to different transcription rates (Figure 7B). Comparing 22G-RNA populations for both piRNA sensors we found that the single intron piRNA sensor accumulated 5-6 fold more 22G-RNAs compared to three-intron piRNA sensor in wild-type animals (Figure S7B). This is consistent with a role of Aquarius/EMB-4 alongside HRDE-1 in nuclear RNAi, including 22G-RNA stability and amplification (Sapetschnig et al., 2015). Most endogenous *C. elegans* genes have, in average, four introns (Michael and Manyuan, 1999). Indeed, we observed that gene de-silencing in both *hrde-1* and *emb-4* mutants correlated with 22G-RNA depletion (Figure S8A-B, yellow quadrant). Furthermore, the negative correlation between 22G-RNA abundance and mRNA levels is stronger when all exons of a gene are targeted by 22G-RNAs (Figure S8A-B, red line vs gray line) when considering *hrde-1* and *emb-4* mutant animals.

**Figure 7.**
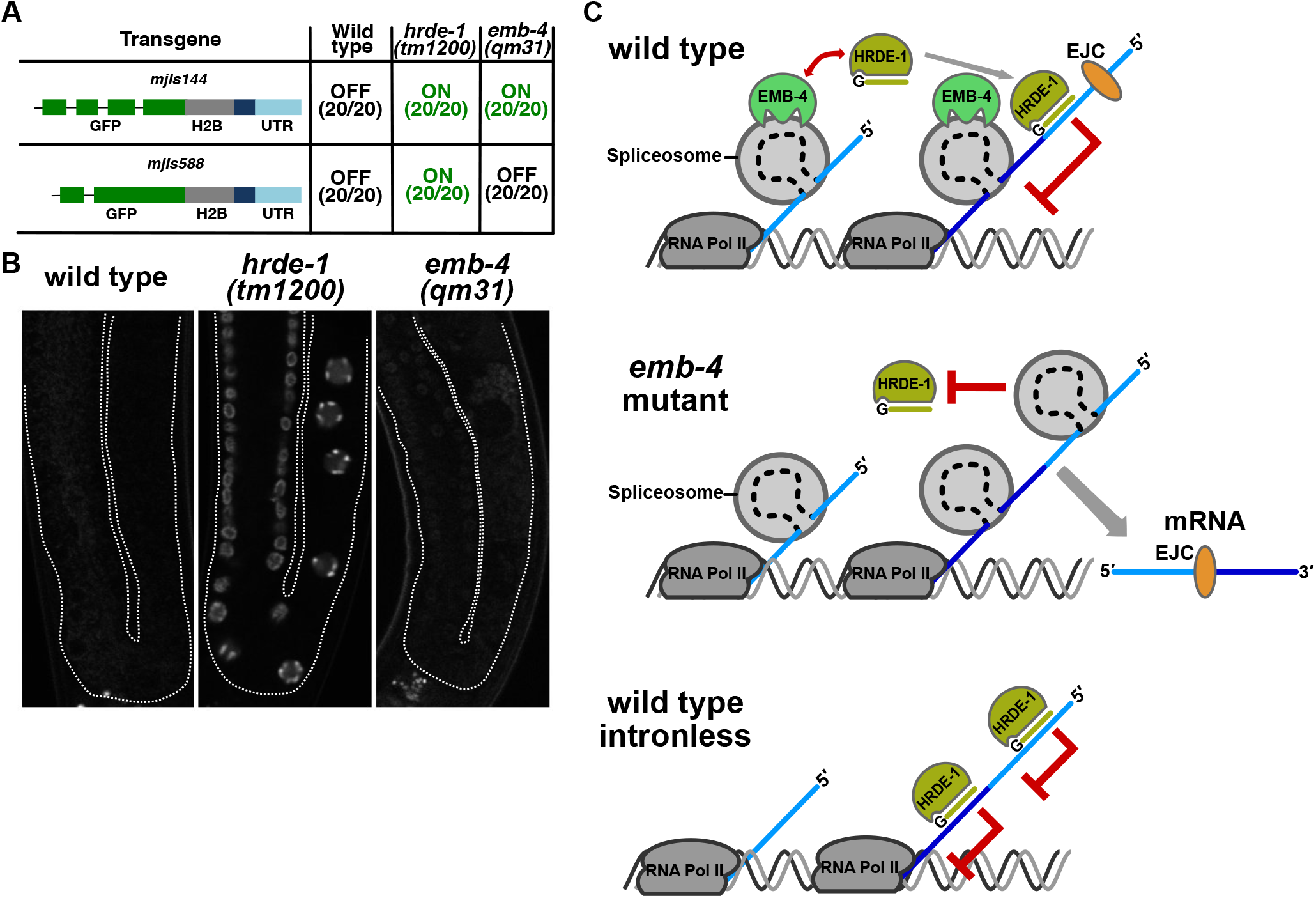
Aquarius/EMB-4 acts to remove intronic barriers to transcriptional gene silencing (A) Silencing of three intron piRNA sensor (*mjIs144*) and single intron piRNA sensor (*mjIs588*) in wild type and mutant animals (20 animals assayed for each condition). (B) Fluorescent microscope images of animals with single intron piRNA sensor (*mjIs588*). (C) Model for Aquarius/EMB-4 function in intron dependent transcriptional gene silencing.

Taken together, our results support a model where the RNA helicase Aquarius/EMB-4 is required to provide the co-transcriptional silencing complex access to nascent transcripts undergoing splicing (Figure 7C). We conclude that pre-mRNA processing is a natural and powerful barrier to co-transcriptional gene silencing.

## DISCUSSION

Here we show that the nuclear Argonaute protein HRDE-1 interacts with spliceosomal and EJC proteins during transcriptional gene silencing in *C. elegans*. Similar interactions have been documented for the *D. melanogaster* nuclear Argonaute PIWI (Czech et al., 2013; Le Thomas et al., 2013),pointing towards a conserved protein complex between nuclear Argonaute proteins involved in co-transcriptional gene silencing and the proteins involved in spliceosome assembly and EJC formation. Several other factors involved in the nuclear RNAi mediated transcriptional gene silencing have been identified in *C. elegans* and *D. melanogaster* using genetic screens (Ashe et al., 2012; Czech et al., 2013; Handler et al., 2013; Lee et al., 2012). These factors include the NRDE factors (NRDE-1/2/4) in *C. elegans* that are required for nuclear RNAi by a yet unknown mechanism (Guang et al., 2008, 2010), as well as several histone methyltransferases and chromatin factors common to both nematodes and flies (Ashe et al., 2012; Shirayama et al., 2012). Much less is known of the events preceding chromatin level changes. In *C. elegans*, HRDE-1 and other nuclear RNAi factors are not only required for transcriptional gene silencing but also affect the biogenesis and/or stability of the 22G-RNAs that are the effector small RNAs for nuclear gene silencing (Sapetschnig et al., 2015). One possible explanation is that in the absence of these factors, interactions between the nuclear Argonaute HRDE-1 and its target RNAs are abolished leading to 22G-RNA destabilisation. It is known that for the miRNA pathway, such Argonaute-target interactions are required to stabilise the associated miRNAs (Chatterjee and Grosshans, 2009; Chatterjee et al., 2011). Considering numerous factors are required for effective HRDE-1 function, it is possible that RNA itself harbors intrinsic features refractory to HRDE-1 targeting.

We identified the spliceosomal RNA helicase Aquarius/EMB-4 as an interactor of HRDE-1 in our proteomics experiments. Aquarius is an intron binding protein that is involved in remodelling during spliceosome and EJC assembly. We show that in *C. elegans* Aquarius/EMB-4 expression is enriched in germ cell nuclei in adult stage animals, which coincides with peak HRDE-1 expression in the germline. Our genetic experiments show that, Aquarius/EMB-4 is required during the initiation step of silencing by HRDE-1 and mutations in the helicase domain of Aquarius/EMB-4 are sufficient to impair its function in transcriptional gene silencing. Absence of Aquarius/EMB-4 leads to transcriptional de-silencing of otherwise silenced genes, transposable elements and the piRNA sensor transgene. Knowing that Aquarius/EMB-4 binds introns led us to investigate the potential role of introns in Aquarius/EMB-4 function.

### Intronic barriers to transcriptional gene silencing

Mammalian Aquarius binds to introns at a certain distance from the intron branch point, in a sequence independent manner (Hirose et al., 2006). Even though the details of such interactions have not been studied in *C. elegans*, EMB-4 has been shown to bind nascent transcripts (Shiimori et al., 2012). Thus, it is reasonable to hypothesise that introns can influence the function of Aquarius/EMB-4 during transcriptional gene silencing. Indeed, when we reduced the number of introns from 3 to 1 in our piRNA sensor transgene, the absence of Aquarius/EMB-4 no longer resulted in de-silencing of the transgene showing that Aquarius/EMB-4 needs intronic sequences to function. Together with our observation that the single intron piRNA sensor accumulates several-fold more 22G-RNAs compared to three-intron piRNA sensor, we propose that introns and/or factors interacting with introns are the inhibitory signals for HRDE-1 silencing.

### Transcription, introns and silencing

Our results add to the growing body of evidence implicating co-transcriptional processes in small RNA mediated gene silencing. In *S. pombe* small RNA mediated coTGS mechanism is required for the silencing of centromeric repeats. Several factors that interact with nascent transcripts and non-essential splicing factors are required for efficient coTGS (Bayne et al., 2008, 2014). In addition, factors influencing the efficiency of RNA polymerase elongation are also important in coTGS mechanism in *S. pombe* (Kowalik et al., 2015). In plants, similar to our results, intron-containing transgenes are protected from nuclear RNAi pathway in comparison to the strongly silenced intronless transgenes (Christie et al., 2011). One hypothesis is that introns or factors interacting with introns are capable of blocking the nuclear RdRPs found in yeast and plants (Dumesic and Madhani, 2013; Vermeersch et al., 2010). In contrast, in the pathogenic yeast *Cryptococcus neoformans*, unspliced introns are a signal for small RNA biogenesis and silencing of genes and transposable elements (Dumesic et al., 2013). Clearly, coTGS mechanisms in different organisms have evolved, one way or the other, to incorporate co-transcriptional processes for the regulation of gene silencing. Unlike in yeast and plants, animals do not possess have nuclear RdRPs, and although *C. elegans* relies on RdRPs for small RNA amplification, this process occurs in the cytoplasm (Phillips et al., 2012). Nuclear processes that lead to coTGS in nematodes, flies and mammals instead rely on the transport of small RNAs from cytoplasm to the nucleus by Argonaute proteins. Our results show that in animal coTGS pathways, introns can pose a barrier to transcriptional gene silencing and we provide evidence that the conserved spliceosomal helicase Aquarius/EMB-4 is required to remove these inhibitory signals.

## AUTHOR CONTRIBUTIONS

A.A. and E.A.M. conceived and designed the study and wrote the manuscript. A.A., K.S., A.N., M.L, R.M, C.J.W performed the experiments. A.A., T.D.D., G.P., M.L., A.B., X.Z., P.M. analysed the data, A.A., R.M., A.N., J.M.C. generated reagents, M.H., J.M.C., A.I.L, E.A.M. provided expertise and feedback.

## ACKNOWLEDGEMENTS

We thank Kay Harnish and Sylviane Moss for high-throughput sequencing support. We thank George Allen and Charles Bradshaw for core computing support. A.B. was supported by a HFSP grant to E.A.M. (RPG0014/2015). This work was supported by Cancer Research UK (C13474/A18583, C6946/A14492), the Wellcome Trust (104640/Z/14/Z, 092096/Z/10/Z) and The European Research Council (ERC, grant 260688). The work of P.M. and X.Z. are supported by NIH grant R01GM113242 and NIH grant R01GM122080. R.M was a Commonwealth Scholar, funded by the UK Government. J.M.C., A.N. and C.J.W were supported by the CIHR (MOP-274660), and the Canada Research Chairs Program.

## ACCESSION NUMBERS

The RNA-sequencing data sets reported in this article have been deposited at EMBL-EBI in ArrayExpress and are accessible through the series accession number E-MTAB-4877. The mass spectrometry proteomics data have been deposited to the ProteomeXchange Consortium via the PRIDE (Vizcaíno et al., 2016) partner repository with the dataset identifier PXD004416. Reviewer account details: Username, password: ILEJacdD.

## EXPERIMENTAL PROCEDURES

### Genetics

*C. elegans* were grown under standard conditions at 20°C unless otherwise indicated. The wild-type strain was var. Bristol N2 (Brenner, 1974).All strains used are listed in Supplementary table 2.

### SILAC Proteomics

Bacterial and nematode growth conditions for SILAC experiments are previously described (Larance et al., 2011).Heavy (R10K8) labelled wild type animals and medium (R6K4) labelled *hrde-1* mutant animals were grown to young adult stage, washed 3× with M9 buffer and lysed in native lysis buffer (10 mM Tris-HCl pH 7.5, 150 mM NaCl, 0.5 mM EDTA, 0.5% NP40, Roche complete protease inhibitor cocktail) by beat beating using zirconia beads and the PreCelys instrument (6,500 rpm, 3×30 s with 30 s intervals at 4°C). The lysate was kept on ice for 30 min and centrifuged for 10 min at 16,000 rcf at 4°C to remove insoluble material. BCA assay (Thermo Scientific) was used to determine protein concentration of the supernatant. 3-8 mg of total protein has been used for immunoprecipitations (IP) with 3 mg of anti-HRDE-1 antibody coupled dynabeads M270 (20 μg antibody / mg beads) for 1 hr at 4°C. Beads were washed 3× with wash buffer (10 mM Tris-HCl pH 7.5, 300 mM NaCl, 0.5 mM EDTA, Roche complete protease inhibitor cocktail) and equal amounts of beads from heavy and medium labelled IPs were mixed together prior to elution at the final wash. Elution was done by heating beads to 70°C for 10 min in LDS loading buffer. The eluted IP was loaded across multiple adjacent lanes (25 μlper lane) on 1 mm, 10-well, 4–12% (w/v) Bis-Tris NuPage gels using MES running buffer according to manufacturer’s instructions but with the addition of 25 mM triscarboxyethylphosphine, and 50 mM N-ethylmaleimide in the LDS sample buffer. After electrophoresis at 150 V for 45 min, SYPRO Ruby staining was performed as per manufacturer’s instructions (Invitrogen).

Protein bands of interest were excised and destained in 1 ml of 50% acetonitrile and 250 mM ammonium bicarbonate at room temperature for 45 min with shaking. The gel slice was dehydrated by incubation in 1 ml of 100% acetonitrile for 10 min at room temperature. All solution was carefully removed prior to the addition of 50 μl MS-grade trypsin (Promega) (12.5 ng/μl) in 100 mM NH4HCO3 and incubation overnight at 37°C. Peptides were extracted by the addition of 0.1 ml of 5% formic acid and incubation at 37°C for 1 hr. Peptides were further extracted by the addition of 0.1 ml of 100% acetonitrile and incubation at 37°C for 1 hr. The gel slice was completely dehydrated by the addition of 0.5 ml of 100% acetonitrile and incubation at 37 °C for 10 min. The entire supernatant was then vacuum-dried.

### LC-MS/MS and analysis of spectra

Using a Thermo Fisher Scientific Ultimate 3000 RSLCnano UHPLC, 15 μl of peptides in 5% (vol/vol) formic acid (final volume ~10 μl) were injected onto an Acclaim PepMap C18 nano-trap column. After washing with 2% (vol/vol) acetonitrile, 0.1% (vol/vol) formic acid, peptides were resolved on a 50 cm × 75 μm C18 EasySpray reverse phase analytical column with integrated emitter over a gradient from 2% acetonitrile to 35% acetonitrile over 220 min with a flow rate of 200 nl/min. The peptides were ionized by electrospray ionization at +2.0 kV. Tandem mass spectrometry analysis was carried out on a Q-Exactive mass spectrometer (Thermo Fisher Scientific) using HCD fragmentation. The data-dependent acquisition method used acquired MS/MS spectra on the top 30 most abundant ions at any one point during the gradient. The RAW data produced by the mass spectrometer were analysed using the MaxQuant quantitative proteomics software package (Cox and Mann, 2008) (http://www.maxquant.org, version 1.3.0.5). The MaxQuant output has also been uploaded to the ProteomeXchange Consortium under the same identifier given above. This version of MaxQuant includes an integrated search engine, Andromeda. Peptide and Protein level identification were both set to a false discovery rate of 1% using a target-decoy based strategy. The database supplied to the search engine for peptide identifications was the combined *C. elegans* and *E. coli* Swissprot and Trembl databases downloaded on the 12th July 2012. The mass tolerance was set to 7 ppm for precursor ions and MS/MS mass tolerance was set at 20 ppm. Enzyme was set to trypsin (cleavage C-terminal to lysine and arginine) with up to 2 missed cleavages. Deamidation of Asn and Gln, oxidation of Met, pyro-Glu (with peptide N-term Gln), phosphorylation of Ser/Thr/Tyr, and protein N-terminal acetylation were set as variable modifications. N-ethylmaleimide on Cys was searched as a fixed modification. The output from MaxQuant provided peptide level data as well as protein group level data. We used the protein groups as defined by the Maxquant package (Cox and Mann, 2008).

### HRDE-1 SILAC IP experimental design, statistical rationale and data analysis

Three biological replicates were performed for SILAC-IP analysis of HRDE-1 and this level of replication was chosen based upon the variance detected in previous experiments using SILAC-IP analysis (Larance et al., 2012).To avoid disregarding low affinity binders, we used a low stringency cutoff such that a protein needed to have a H/M SILAC ratio >1 in two out of three biological replicates in our data analysis with MaxQuant to eliminate non-specific binding proteins. This yields excellent removal of environmental contaminants (keratins, trypsin, antibody, etc.) that do not incorporate stable isotopes.

### HRDE-1 / EMB-4 co-immunoprecipitations and western blot analysis

For HRDE-1 immunoprecipitations, animals were harvested 24 hours post-L4 stage in 30 mM HEPES, 100 mM potassium acetate, 2 mM magnesium acetate and 10% glycerol (DROSO buffer). To lyse the animals, samples were snap-frozen in liquid nitrogen as droplets and grinded to powder. *C. elegans* powder was then re-suspended in DROSO buffer supplemented with 0.01% NP-40, 2 mM DTT and further lysed by bead-beating. Lysates were subsequently cleared by centrifugation. 2 mg of proteins were used per IP at 4 mg/ml. 10 μg antibody (normal IgG: SantaCruz Biotech, sc-2027; anti-HRDE-1: Genomic Antibody Tech, custom (Kamminga et al., 2012))was added to the lysates and incubated overnight at 4°C with rotation. 30 μl of proteinA/G-agarose beads (SantaCruz Biotech, sc-2003) were incubated for 2 hrs the next morning. Immunoprecipitates were washed 4 times with DROSO buffer and boiled with 2× sample buffer for 5 min to elute. Samples were then analysed by western blotting (Anti-Ollas: Novusbio, NMP1-06713).

For EMB-4 immunoprecipitations, 750 µl of synchronized gravid adults were dounced using a metal wheaton dounce in DROSO ‘complete’ buffer (30 mM Hepes, 100 mM potassium Acetate, 2 mM Magnesium Acetate, 0.1% NP40/Igepal, 2 mM DTT, 1 tablet/5mls Protease inhibitor (Roche), 1:100 Sigma Phosphatase Inhibitor 2, 1:100 Sigma Phosphatase Inhibitor 3.) until the worms and the embryos were no longer visible. Lysate was cleared by centrifugation for 10min at 13,000xg in a pre-cooled centrifuge (4°C). The concentration of the supernatant (total worm protein) was determined by Lowry assay using Bio-rad Lowry assay kit. Each IP was performed from 5 mg protein. Lysate was pre-cleared with 25 µl protein A/G agarose bead slurry (Santa Cruz Biotech, beads are equilibrated in DROSO complete buffer prior to use) for one hour at 4°C on a rotator. 5 µg (anti-Flag, Sigma Aldrich) or 50 µl of EMB-4 antibody (specificity of EMB-4 antibodies are shown in Figure S1E) or buffer alone (no antibody control) was added to each IP sample and incubated for two hours on a rotator at 4°C. Immune complexes were recovered using 50µl of a 50% slurry of Protein-A/G agarose beads (Santa Cruz Biotechnology) and washed 6x5min at 4°C with DROSO buffer. Protein was eluted from beads and denatured by incubation in Thermofisher 2x LDS sample buffer for 10min at 70°C. Input samples were prepared from the same lysate at a concentration of 2ug/ul using Thermofisher 2xLDS sample buffer and reducing agent. Proteins were resolved by SDS-PAGE on Criterion Precast gradient gels (4-15%, Biorad) and transferred to Hybond-C membrane(Amersham Biosciences). The membrane was incubated overnight at 4°C with either: (i) affinity purified anti-EMB-4 (1:200), or anti-FLAG (Sigma Aldrich, 1:1000) in PBST-5% milk solution (137 mM NaCl, 10 mM Phosphate, 2.7 mM KCl, pH 7.4, and 5% [w/v] dried milk). The membrane was incubated 2 h at room temperature with anti-mouse HRP-conjugated secondary antibody (Jackson Immunoresearch) diluted 1:1,000 in PBST and then visualized by Luminata Forte Western HRP substrate.

### qRT-PCR analysis of GFP expression

cDNA is synthesised from purified and DNase treated total RNA using the SuperScript II enzyme as described in the manual. qRT-PCR reactions are performed using Applied Biosystem Power SYBR Green Master Mix using primer sequences to amplify GFP sequence (primer 1: 5′-TCTGTCAGTGGAGAGGGTGA-3′, primer2: 5′-TTTAAACTTACCCATGGAACAGG-3′).

### qRT-PCR analysis of *emb-4* expression

cDNA was generated from 1mg of *C. elegans* total RNA using random hexamers with Superscript III Reverse Transcriptase (Invitrogen). qRT-PCR was performed using Applied Biosystems SYBR Green PCR Master mix with primers for *emb-4* (primer 1: 5′-TTCGTCCCCTGTTCCATATC-3′, primer 2: 5′-ATCGGCTTCTGGCCTAAAAT-3′) and for *act-3* (primer 1: 5′-CCAAGAGAGGTATCCTTACCCTCAA-3′, primer 2: 5′-AAGCTCATTGTAGAAGGTGTGATGC-3′).

### Western blot analysis of EMB-4 expression

Proteins were resolved by SDS-PAGE on Criterion Precast gradient gels (4-15%, Biorad). and transferred to Hybond-C membrane (Amersham Biosciences). The membrane was incubated overnight at 4°C with either: (i) affinity purified anti-EMB-4, or anti alpha-tubulin (Accurate Chemical) antibodies diluted to 1:2000, in PBST-5% milk solution (137 mM NaCl, 10 mM Phosphate, 2.7 mM KCl, pH 7.4, and 5% [w/v] dried milk). The membrane was incubated 1 h at room temperature with HRP-conjugated secondary antibodies (Jackson Immunoresearch) diluted to 1:5,000 in PBST and then visualized by Western Lightning ECL Kit from Perkin Elmer. Images were collected on a LAS-3000 Intelligent Dark-Box (Fujifilm).

### Immuno-staining of *C. elegans* gonads and embryos

Gonads and embryos were excised from gravid adult worms in 1x PBS on poly-L-lysine coated slides, frozen and cracked on dry ice for longer than 10 minutes, and fixed at –20°C for 5 min in each of the following (15 minutes total) respectively; 100% methanol, 50% methanol/50% acetone, and 100% acetone. All sample incubations were performed in a humid chamber. Samples were washed 2x 5min with 1xPBS, then 2x 5 mins with 1xPBS / 0.1% Tween-20. Samples are then blocked for one hour in 1xPBS/0.1% Tween-20 / 3%BSA (PBST+BSA) at room temperature, and then incubated with primary antibody (1:500) overnight at 4°C. Slides were washed 3x for 10 minutes with PBST, and then incubated for 1 hour in PBST+BSA. Secondary antibodies were from Jackson Immunoresearch and Molecular Probes. Incubations with anti-mouse secondary antibodies were performed for one hour in PBST+BSA at room temperature. Slides were washed 3x for ten minutes in PBST, 3x for 5 minutes in PBS and then incubated with DAPI (1:2500) for 10 minutes at room temperature. Finally, slides were washed in PBS 3x for 5 minutes then mounted in Vectashield (Vector Labs). All images were collected using Nikon Ti-S inverted microscope with NIS Element and AR software.

### RNA sequencing

Synchronised animals were grown to young adult stage at 20°C on HB101 seeded NGM plates. Animals were harvested and washed 3X in M9 buffer. Settled animals were mixed with Trisure reagent, bead beaten as described in proteomics experiments above and total RNA was isolated by a chloroform extraction.

For total RNA sequencing, Illumina Ribozero kit was used to remove ribosomal RNA from 1 μg of total RNA prior to library preparation. RNA sequencing libraries were prepared using NEB Next Ultra library preparation kit. Small RNA sequencing performed by treating 5 μg of total RNA with Epicentre 5′ polyphosphatase to remove the 5′ triphosphate from 22G-RNAs. After treatment, RNA is purified by phenol/chloroform extraction and 1 μg of RNA is used to prepare small RNA libraries using Illumina TruSeq small RNA library preparation kit. Ribosomal depleted RNA and small RNA libraries are sequenced using Illumina HiSeq 1500 platform.

### RNA sequence analysis

The ce10/WS220 genome fasta file was obtained from the WormBase ftp server. Sequences for the piRNA sensor transgene and the piRNA sensor transgene with one intronwere added as separate chromosomes when required. A GTF file containing annotations for genes and pseudogene for version ce10/WS220 of the *C. elegans* genome were downloaded from the UCSC table browser. To prevent multiple counts per read in the case of overlapping features, only the longest isoform of each gene was included in the analysis. A GTF file containing annotations for transposable elements was generated by running RepeatMasker version open-4.0.5 in sensitive mode, run with rmblastn version 2.2.27+ using RepeatMasker database version 20140131, against the ce10/WS220 genome fasta file. “Simple_repeat” and “Low_complexity” annotations were excluded from the analysis. Raw fastq small RNA sequencing files were processed by removing the Illumina TruSeq adaptor sequence using cutadapt v1.9, with parameters “--minimum-length 18 --discard-untrimmed -a TGGAATTCTCGGGTGCCAAGG”. Raw fastq RNA sequencing files were processed by removing the NEBNext adaptor sequence using cutadapt v1.9, with parameters “-a AGATCGGAAGAGCACACGTCTGAACTCCAGTCAC”. Adaptor-trimmed small RNA sequencing reads were aligned to the ce10/WS220 *C. elegans* genome using STAR v2.5.1b, with parameters “--outFilterMultimapNmax 50 --winAnchorMultimapNmax 50 --outFilterMismatchNmax 0 --limitBAMsortRAM 31000000000 --alignIntronMax 1 --alignEndsType EndToEnd --outSAMtype BAM SortedByCoordinate --runThreadN 6 --outBAMsortingThreadN 6 --readFilesCommand ‘gunzip -c’”;. Adaptor-trimmed RNA sequencing reads were aligned to the ce10/WS220 *C. elegans* genome using STAR v2.5.1b, with parameters “--outFilterMultimapNmax 5000 --winAnchorMultimapNmax 10000 --outFilterMismatchNmax 2 --alignEndsType EndToEnd --outSAMtype BAM Unsorted --runThreadN --readFilesCommand ‘gunzip -c’”. Aligned RNA sequencing reads were sorted and indexed using samtools v1.3. Counts against the annotations in the GTF files were generated with featureCounts v1.5.0-p1, with parameters “-T 6 -M --fraction”. Normalised counts, variance-stabilised counts, fold change values, and adjusted p-values were obtained using DESeq2 v3.2.2, called through a custom script.

We also used exon-intron split analysis (EISA) (Gaidatzis et al., 2015) to characterize the gene expression changes detected between *hrde-1* or *emb-4* null and wild-type strains. Both exonic and intronic read counts were quantified using FeatureCounts (Liao et al., 2014). When using EISA, we processed the counts and the annotation files by following the procedures described by (Gaidatzis et al., 2015).

We calculated the 22G-RNA density using previously published small RNA sequencing data obtained from HRDE-1 immunoprecipitations in wild type and mutant animals normalised to library size (Sapetschnig et al., 2015).We used a cut-off of 22G-RNA reads in wild type / *hrde-1* mutant control ≥ 4 for filtering out 22G-RNA reads that were unspecifically binding to anti-HRDE-1 antibody. We then used the following calculation 22G-RNA density=# of 22G-RNA reads in HRDE-1 IP of gene A / RPKM of the gene A.

We carried out exon level sRNA differential expression by filtering out genes that have zero mapped reads in all samples and normalising the samples by sample size using the Median Ratio Method (Anders and Huber, 2010) implemented in the R package DESeq2 to adjust for factors like the coverage and sampling depth. Next, we log transformed the exon read counts for each gene, performed a two-sample t-test on each exon independently and adjusted the p-values of testing results by false discovery rate using the Benjamini & Hochberg method (Benjamini and Hochberg, 1995).We also performed an analysis of sRNA dependent gene regulation on genes with or without isoforms: We took genes that show log fold change of mRNA greater than 1 and log fold change of sRNA less than -0.6 and analysed the number of genes with annotated isoforms and number of genes with no isoforms. The isoform annotation is based on *C. elegans* genome release CE10 (WS220). We used Chi-squared test to test if the above gene set has different proportion of genes with isoforms to all genes or germline specific sRNA target genes (Gu et al., 2009b).

### Transgenic animals

The *mjIs588* allele was generated by removing the introns two and three from the GFP sequence in the plasmid pEM975 that is used to generate the *mjIs144* allele. New plasmid is inserted on Chr II using the previously described MosSCI method (Frøkjær-Jensen et al., 2008) into the same location as in *mjIs144* allele. *mjSi92* allele is generated by CRISPR tagging of endogenous *emb-4* N-term with the OLLAS epitope sequence using CRISPR gRNA (5′-CAAGAAGCCGTGGTGACTCG-3′) and the repair template plasmid pEM2058 using Cas9 protein and RNA injections as described in Paix et al. (Paix et al., 2015).

### Structural alignment of AQR and EMB-4

EMB-4 structure was determined by PHYRE-2 online prediction tool. Images were generated in Pymol (The PyMOL Molecular Graphics System, Version 1.8 Schrödinger, LLC) and mutagenesis was performed in Chimera (Yang et al., 2012).

## REFERENCES

Akay, A., Craig, A., Lehrbach, N., Larance, M., Pourkarimi, E., Wright, J.E., Lamond, A., Miska, E., and Gartner, A. (2013). RNA-binding protein GLD-1/quaking genetically interacts with the mir-35 and the let-7 miRNA pathways in Caenorhabditis elegans. Open Biol. 3, 130151.

de Albuquerque, B.F.M., Placentino, M., and Ketting, R.F. (2015). Maternal piRNAs Are Essential for Germline Development following De Novo Establishment of Endo-siRNAs in Caenorhabditis elegans. Dev. Cell 34, 448–456.

Anders, S., and Huber, W. (2010). Differential expression analysis for sequence count data. Genome Biol. 11, R106.

Aravin, A.A., Sachidanandam, R., Bourc’his, D., Schaefer, C., Pezic, D., Toth, K.F., Bestor, T., and Hannon, G.J. (2008). A piRNA pathway primed by individual transposons is linked to de novo DNA methylation in mice. Mol. Cell 31, 785–799.

Ashe, A., Sapetschnig, A., Weick, E.-M., Mitchell, J., Bagijn, M.P., Cording, A.C., Doebley, A.-L., Goldstein, L.D., Lehrbach, N.J., Le Pen, J., et al. (2012). piRNAs can trigger a multigenerational epigenetic memory in the germline of C. elegans. Cell 150, 88–99.

Bagijn, M.P., Goldstein, L.D., Sapetschnig, A., Weick, E.-M., Bouasker, S., Lehrbach, N.J., Simard, M.J., and Miska, E.A. (2012). Function, targets, and evolution of Caenorhabditis elegans piRNAs. Science 337, 574–578.

Batista, P.J., Ruby, J.G., Claycomb, J.M., Chiang, R., Fahlgren, N., Kasschau, K.D., Chaves, D.A., Gu, W., Vasale, J.J., Duan, S., et al. (2008). PRG-1 and 21U-RNAs interact to form the piRNA complex required for fertility in C. elegans. Mol. Cell 31, 67–78.

Bayne, E.H., Portoso, M., Kagansky, A., Kos-Braun, I.C., Urano, T., Ekwall, K., Alves, F., Rappsilber, J., and Allshire, R.C. (2008). Splicing factors facilitate RNAi-directed silencing in fission yeast. Science 322, 602–606.

Bayne, E.H., Bijos, D.A., White, S.A., de Lima Alves, F., Rappsilber, J., and Allshire, R.C. (2014). A systematic genetic screen identifies new factors influencing centromeric heterochromatin integrity in fission yeast. Genome Biol. 15, 481.

Benjamini, Y., and Hochberg, Y. (1995). Controlling the False Discovery Rate: A Practical and Powerful Approach to Multiple Testing. J. R. Stat. Soc. Series B Stat. Methodol. 57, 289–300.

Brennecke, J., Aravin, A.A., Stark, A., Dus, M., Kellis, M., Sachidanandam, R., and Hannon, G.J. (2007). Discrete small RNA-generating loci as master regulators of transposon activity in Drosophila. Cell 128, 1089–1103.

Brenner, S. (1974). The genetics of Caenorhabditis elegans. Genetics 77, 71–94.

Buckley, B.A., Burkhart, K.B., Gu, S.G., Spracklin, G., Kershner, A., Fritz, H., Kimble, J., Fire, A., and Kennedy, S. (2012). A nuclear Argonaute promotes multigenerational epigenetic inheritance and germline immortality. Nature 489, 447–451.

Burkhart, K.B., Guang, S., Buckley, B.A., Wong, L., Bochner, A.F., and Kennedy, S. (2011). A Pre-mRNA–Associating Factor Links Endogenous siRNAs to Chromatin Regulation. PLoS Genet. 7, e1002249.

Carmell, M.A., Girard, A., van de Kant, H.J.G., Bourc’his, D., Bestor, T.H., de Rooij, D.G., and Hannon, G.J. (2007). MIWI2 is essential for spermatogenesis and repression of transposons in the mouse male germline. Dev. Cell 12, 503–514.

Carrillo Oesterreich, F., Herzel, L., Straube, K., Hujer, K., Howard, J., and Neugebauer, K.M. (2016). Splicing of Nascent RNA Coincides with Intron Exit from RNA Polymerase II. Cell 165, 372–381.

Chakrabarti, S., Jayachandran, U., Bonneau, F., Fiorini, F., Basquin, C., Domcke, S., Le Hir, H., and Conti, E. (2011). Molecular mechanisms for the RNA-dependent ATPase activity of Upf1 and its regulation by Upf2. Mol. Cell 41, 693–703.

Chatterjee, S., and Grosshans, H. (2009). Active turnover modulates mature microRNA activity in Caenorhabditis elegans. Nature 461, 546–549.

Chatterjee, S., Fasler, M., Büssing, I., and Grosshans, H. (2011). Target-mediated protection of endogenous microRNAs in C. elegans. Dev. Cell 20, 388–396.

Checchi, P.M., and Kelly, W.G. (2006). emb-4 is a conserved gene required for efficient germline-specific chromatin remodeling during Caenorhabditis elegans embryogenesis. Genetics 174, 1895–1906.

Cheng, Z., Muhlrad, D., Lim, M.K., Parker, R., and Song, H. (2007). Structural and functional insights into the human Upf1 helicase core. EMBO J. 26, 253–264.

Christie, M., Croft, L.J., and Carroll, B.J. (2011). Intron splicing suppresses RNA silencing in Arabidopsis. Plant J. 68, 159–167.

Claycomb, J.M., Batista, P.J., Pang, K.M., Gu, W., Vasale, J.J., van Wolfswinkel, J.C., Chaves, D.A., Shirayama, M., Mitani, S., Ketting, R.F., et al. (2009). The Argonaute CSR-1 and its 22G-RNA cofactors are required for holocentric chromosome segregation. Cell 139, 123–134.

Conine, C.C., Batista, P.J., Gu, W., Claycomb, J.M., Chaves, D.A., Shirayama, M., and Mello, C.C. (2010). Argonautes ALG-3 and ALG-4 are required for spermatogenesis-specific 26G-RNAs and thermotolerant sperm in Caenorhabditis elegans. Proc. Natl. Acad. Sci. U. S. A. 107, 3588–3593.

Cowley, M., and Oakey, R.J. (2013). Transposable elements re-wire and fine-tune the transcriptome. PLoS Genet. 9, e1003234.

Cox, J., and Mann, M. (2008). MaxQuant enables high peptide identification rates, individualized ppb-range mass accuracies and proteome-wide protein quantification. Nat. Biotechnol. 26, 1367–1372.

Cox, D.N., Chao, A., Baker, J., Chang, L., Qiao, D., and Lin, H. (1998). A novel class of evolutionarily conserved genes defined by piwi are essential for stem cell self-renewal. Genes Dev. 12, 3715–3727.

Czech, B., Preall, J.B., McGinn, J., and Hannon, G.J. (2013). A Transcriptome-wide RNAi Screen in the Drosophila Ovary Reveals Factors of the Germline piRNA Pathway. Mol. Cell 50, 1–13.

Das, P.P., Bagijn, M.P., Goldstein, L.D., Woolford, J.R., Lehrbach, N.J., Sapetschnig, A., Buhecha, H.R., Gilchrist, M.J., Howe, K.L., Stark, R., et al. (2008). Piwi and piRNAs act upstream of an endogenous siRNA pathway to suppress Tc3 transposon mobility in the Caenorhabditis elegans germline. Mol. Cell 31, 79–90.

De, I., Bessonov, S., Hofele, R., dos Santos, K., Will, C.L., Urlaub, H., Lührmann, R., and Pena, V. (2015). The RNA helicase Aquarius exhibits structural adaptations mediating its recruitment to spliceosomes. Nat. Struct. Mol. Biol. 22, 138–144.

Drinnenberg, I.A., Fink, G.R., and Bartel, D.P. (2011). Compatibility with killer explains the rise of RNAi-deficient fungi. Science 333, 1592.

Dumesic, P.A., and Madhani, H.D. (2013). The spliceosome as a transposon sensor. RNA Biol. 10, 1653–1660.

Dumesic, P.A., Natarajan, P., Chen, C., Drinnenberg, I.A., Schiller, B.J., Thompson, J., Moresco, J.J., Yates, J.R., 3rd, Bartel, D.P., and Madhani, H.D. (2013). Stalled spliceosomes are a signal for RNAi-mediated genome defense. Cell 152, 957–968.

Fire, A., Harrison, S.W., and Dixon, D. (1990). A modular set of lacZ fusion vectors for studying gene expression in Caenorhabditis elegans. Gene 93, 189–198.

Fire, A., Xu, S., Montgomery, M.K., Kostas, S.A., Driver, S.E., and Mello, C.C. (1998). Potent and specific genetic interference by double-stranded RNA in Caenorhabditis elegans. Nature 391, 806–811.

Fredens, J., Engholm-Keller, K., Giessing, A., Pultz, D., Larsen, M.R., Højrup, P., Møller-Jensen, J., and Færgeman, N.J. (2011). Quantitative proteomics by amino acid labeling in C. elegans. Nat. Methods 8, 845–847.

Frøkjær-Jensen, C., Christian, F.-J., Wayne Davis, M., Hopkins, C.E., Newman, B.J., Thummel, J.M., Søren-Peter, O., Morten, G., and Jorgensen, E.M. (2008). Single-copy insertion of transgenes in Caenorhabditis elegans. Nat. Genet. 40, 1375–1383.

Gaidatzis, D., Burger, L., Florescu, M., and Stadler, M.B. (2015). Analysis of intronic and exonic reads in RNA-seq data characterizes transcriptional and post-transcriptional regulation. Nat. Biotechnol. 33, 722–729.

Gerson-Gurwitz, A., Wang, S., Sathe, S., Green, R., Yeo, G.W., Oegema, K., and Desai, A. (2016). A Small RNA-Catalytic Argonaute Pathway Tunes Germline Transcript Levels to Ensure Embryonic Divisions. Cell 165, 396–409.

Gonatopoulos-Pournatzis, T., and Cowling, V.H. (2014). Cap-binding complex (CBC). Biochem. J 457, 231–242.

Gu, W., Shirayama, M., Conte, D., Jr., Vasale, J., Batista, P.J., Claycomb, J.M., Moresco, J.J., Youngman, E.M., Keys, J., Stoltz, M.J., et al. (2009a). Distinct Argonaute-Mediated 22G-RNA Pathways Direct Genome Surveillance in the C. elegans Germline. Mol. Cell 36, 231–244.

Gu, W., Shirayama, M., Conte, D., Jr., Vasale, J., Batista, P.J., Claycomb, J.M., Moresco, J.J., Youngman, E.M., Keys, J., Stoltz, M.J., et al. (2009b). Distinct Argonaute-Mediated 22G-RNA Pathways Direct Genome Surveillance in the C. elegans Germline. Mol. Cell 36, 231–244.

Guang, S., Bochner, A.F., Pavelec, D.M., Burkhart, K.B., Harding, S., Lachowiec, J., and Kennedy, S. (2008). An Argonaute Transports siRNAs from the Cytoplasm to the Nucleus. Science 321, 537–541.

Guang, S., Bochner, A.F., Burkhart, K.B., Burton, N., Pavelec, D.M., and Kennedy, S. (2010). Small regulatory RNAs inhibit RNA polymerase II during the elongation phase of transcription. Nature 465, 1097–1101.

Handler, D., Meixner, K., Pizka, M., Lauss, K., Schmied, C., Gruber, F.S., and Brennecke, J. (2013). The genetic makeup of the Drosophila piRNA pathway. Mol. Cell 50, 762–777.

Hirose, T., Ideue, T., Nagai, M., Hagiwara, M., Shu, M.-D., and Steitz, J.A. (2006). A spliceosomal intron binding protein, IBP160, links position-dependent assembly of intron-encoded box C/D snoRNP to pre-mRNA splicing. Mol. Cell 23, 673–684.

Holoch, D., and Moazed, D. (2015). RNA-mediated epigenetic regulation of gene expression. Nat. Rev. Genet. 16, 71–84.

Jobson, M.A., Jordan, J.M., Sandrof, M.A., Hibshman, J.D., Lennox, A.L., and Baugh, L.R. (2015). Transgenerational Effects of Early Life Starvation on Growth, Reproduction, and Stress Resistance in Caenorhabditis elegans. Genetics 201, 201–212.

Kamminga, L.M., van Wolfswinkel, J.C., Luteijn, M.J., Kaaij, L.J.T., Bagijn, M.P., Sapetschnig, A., Miska, E.A., Berezikov, E., and Ketting, R.F. (2012). Differential impact of the HEN1 homolog HENN-1 on 21U and 26G RNAs in the germline of Caenorhabditis elegans. PLoS Genet. 8, e1002702.

Katic, I., Katic, I., Greenwald, I., and Greenwald, I. (2006). EMB-4: a predicted ATPase that facilitates lin-12 activity in Caenorhabditis elegans. Genetics 174, 1907–1915.

Kowalik, K.M., Shimada, Y., Flury, V., Stadler, M.B., Batki, J., and Bühler, M. (2015). The Paf1 complex represses small-RNA-mediated epigenetic gene silencing. Nature 520, 248–252.

Kuramochi-Miyagawa, S., Watanabe, T., Gotoh, K., Totoki, Y., Toyoda, A., Ikawa, M., Asada, N., Kojima, K., Yamaguchi, Y., Ijiri, T.W., et al. (2008). DNA methylation of retrotransposon genes is regulated by Piwi family members MILI and MIWI2 in murine fetal testes. Genes Dev. 22, 908–917.

Larance, M., Bailly, A.P., Pourkarimi, E., Hay, R.T., Buchanan, G., Coulthurst, S., Xirodimas, D.P., Gartner, A., and Lamond, A.I. (2011). Stable-isotope labeling with amino acids in nematodes. Nat. Methods 8, 849–851.

Larance, M., Kirkwood, K.J., Xirodimas, D.P., Lundberg, E., Uhlen, M., and Lamond, A.I. (2012). Characterization of MRFAP1 turnover and interactions downstream of the NEDD8 pathway. Mol. Cell. Proteomics 11, M111.014407.

Lee, H.-C., Gu, W., Shirayama, M., Youngman, E., Conte, D., and Mello, C.C. (2012). C. elegans piRNAs Mediate the Genome-wide Surveillance of Germline Transcripts. Cell 150, 78–87.

Le Hir, H., Saulière, J., and Wang, Z. (2016). The exon junction complex as a node of post-transcriptional networks. Nat. Rev. Mol. Cell Biol. 17, 41–54.

Le Thomas, A., Rogers, A.K., Webster, A., Marinov, G.K., Liao, S.E., Perkins, E.M., Hur, J.K., Aravin, A.A., and Tóth, K.F. (2013). Piwi induces piRNA-guided transcriptional silencing and establishment of a repressive chromatin state. Genes Dev. 27, 390–399.

Liao, Y., Smyth, G.K., and Shi, W. (2014). featureCounts: an efficient general purpose program for assigning sequence reads to genomic features. Bioinformatics 30, 923–930.

Luteijn, M.J., van Bergeijk, P., Kaaij, L.J.T., Almeida, M.V., Roovers, E.F., Berezikov, E., and Ketting, R.F. (2012). Extremely stable Piwi-induced gene silencing in Caenorhabditis elegans. EMBO J. 31, 3422–3430.

Michael, D., and Manyuan, L. (1999). Intron—exon structures of eukaryotic model organisms. Nucleic Acids Res. 27, 3219–3228.

Ni, J.Z., Chen, E., and Gu, S.G. (2014). Complex coding of endogenous siRNA, transcriptional silencing and H3K9 methylation on native targets of germline nuclear RNAi in C. elegans. BMC Genomics 15, 1157.

Ni, J.Z., Kalinava, N., Chen, E., Huang, A., Trinh, T., and Gu, S.G. (2016). A transgenerational role of the germline nuclear RNAi pathway in repressing heat stress-induced transcriptional activation in C. elegans. Epigenetics Chromatin 9, 3.

Nilsen, T.W. (2003). The spliceosome: the most complex macromolecular machine in the cell? Bioessays 25, 1147–1149.

Ong, S.-E., Blagoev, B., Kratchmarova, I., Kristensen, D.B., Steen, H., Pandey, A., and Mann, M. (2002). Stable isotope labeling by amino acids in cell culture, SILAC, as a simple and accurate approach to expression proteomics. Mol. Cell. Proteomics 1, 376–386.

Paix, A., Folkmann, A., Rasoloson, D., and Seydoux, G. (2015). High Efficiency, Homology-Directed Genome Editing in Caenorhabditis elegans Using CRISPR-Cas9 Ribonucleoprotein Complexes. Genetics 201, 47–54.

Pak, J., and Fire, A. (2007). Distinct populations of primary and secondary effectors during RNAi in C. elegans. Science 315, 241–244.

Park, S.H., Cheong, C., Idoyaga, J., Kim, J.Y., Choi, J.-H., Do, Y., Lee, H., Jo, J.H., Oh, Y.-S., Im, W., et al. (2008). Generation and application of new rat monoclonal antibodies against synthetic FLAG and OLLAS tags for improved immunodetection. J. Immunol. Methods 331, 27–38.

Phillips, C.M., Montgomery, T.A., Breen, P.C., and Ruvkun, G. (2012). MUT-16 promotes formation of perinuclear Mutator foci required for RNA silencing in the C. elegans germline. Genes Dev. 26, 1433–1444.

Phillips, C.M., Brown, K.C., Montgomery, B.E., Ruvkun, G., and Montgomery, T.A. (2015). piRNAs and piRNA-Dependent siRNAs Protect Conserved and Essential C. elegans Genes from Misrouting into the RNAi Pathway. Dev. Cell 34, 457–465.

Rebollo, R., Romanish, M.T., and Mager, D.L. (2012). Transposable elements: an abundant and natural source of regulatory sequences for host genes. Annu. Rev. Genet. 46, 21–42.

Rechavi, O., Houri-Ze’evi, L., Anava, S., Goh, W.-S.S., Kerk, S.Y., Hannon, G.J., and Hobert, O. (2014). Starvation-induced transgenerational inheritance of small RNAs in C. elegans. Cell 158, 277–287.

Sapetschnig, A., Sarkies, P., Lehrbach, N.J., and Miska, E.A. (2015). Tertiary siRNAs Mediate Paramutation in C. elegans. PLoS Genet. 11, e1005078.

Shatkin, A.J., and Manley, J.L. (2000). The ends of the affair: capping and polyadenylation. Nat. Struct. Biol. 7, 838–842.

Shiimori, M., Inoue, K., and Sakamoto, H. (2012). A Specific Set of Exon Junction Complex Subunits Is Required for the Nuclear Retention of Unspliced RNAs in Caenorhabditis elegans. Mol. Cell. Biol. 33, 444–456.

Shirayama, M., Seth, M., Lee, H.-C., Gu, W., Ishidate, T., Conte, D., and Mello, C.C. (2012). piRNAs Initiate an Epigenetic Memory of Nonself RNA in the C. elegans Germline. Cell 150, 65–77.

Simon, M., Sarkies, P., Ikegami, K., Doebley, A.-L., Goldstein, L.D., Mitchell, J., Sakaguchi, A., Miska, E.A., and Ahmed, S. (2014). Reduced insulin/IGF-1 signaling restores germ cell immortality to Caenorhabditis elegans Piwi mutants. Cell Rep. 7, 762–773.

Slotkin, R.K., and Martienssen, R. (2007). Transposable elements and the epigenetic regulation of the genome. Nat. Rev. Genet. 8, 272–285.

Szklarczyk, D., Franceschini, A., Wyder, S., Forslund, K., Heller, D., Huerta-Cepas, J., Simonovic, M., Roth, A., Santos, A., Tsafou, K.P., et al. (2015). STRING v10: protein-protein interaction networks, integrated over the tree of life. Nucleic Acids Res. 43, D447–D452.

Uhlén, M., Fagerberg, L., Hallström, B.M., Lindskog, C., Oksvold, P., Mardinoglu, A., Sivertsson, Å., Kampf, C., Sjöstedt, E., Asplund, A., et al. (2015). Proteomics. Tissue-based map of the human proteome. Science 347, 1260419.

UniProt Consortium (2015). UniProt: a hub for protein information. Nucleic Acids Res. 43, D204–D212.

Vasale, J.J., Gu, W., Thivierge, C., Batista, P.J., Claycomb, J.M., Youngman, E.M., Duchaine, T.F., Mello, C.C., and Conte, D. (2010). Sequential rounds of RNA-dependent RNA transcription drive endogenous small-RNA biogenesis in the ERGO-1/Argonaute pathway. Proc. Natl. Acad. Sci. U. S. A.

Vermeersch, L., De Winne, N., and Depicker, A. (2010). Introns reduce transitivity proportionally to their length, suggesting that silencing spreads along the pre-mRNA. Plant J. 64, 392–401.

Vizcaíno, J.A., Csordas, A., del-Toro, N., Dianes, J.A., Griss, J., Lavidas, I., Mayer, G., Perez-Riverol, Y., Reisinger, F., Ternent, T., et al. (2016). 2016 update of the PRIDE database and its related tools. Nucleic Acids Res. 44, D447–D456.

Wahl, M.C., and Lührmann, R. (2015). SnapShot: Spliceosome Dynamics I. Cell 161, 1474–e1.

Wang, G., and Reinke, V. (2008). A C. elegans Piwi, PRG-1, regulates 21U-RNAs during spermatogenesis. Curr. Biol. 18, 861–867.

Weick, E.-M., and Miska, E.A. (2014). piRNAs: from biogenesis to function. Development 141, 3458–3471.

Weick, E.-M., Sarkies, P., Silva, N., Chen, R.A., Moss, S.M.M., Cording, A.C., Ahringer, J., Martinez-Perez, E., and Miska, E.A. (2014). PRDE-1 is a nuclear factor essential for the biogenesis of Ruby motif-dependent piRNAs in C. elegans. Genes Dev. 28, 783–796.

Yang, Z., Lasker, K., Schneidman-Duhovny, D., Webb, B., Huang, C.C., Pettersen, E.F., Goddard, T.D., Meng, E.C., Sali, A., and Ferrin, T.E. (2012). UCSF Chimera, MODELLER, and IMP: an integrated modeling system. J. Struct. Biol. 179, 269–278.

